# PAK4 promotes vertex remodeling to maintain epithelial integrity and barrier function

**DOI:** 10.1101/2025.10.02.680170

**Authors:** Babli Adhikary, Alexander Chang, Atsuko Y. Higashi, Hideki Chiba, Ann L. Miller, Tomohito Higashi

## Abstract

Cell-cell junctions are essential for epithelial integrity and barrier function, but the mechanisms regulating their remodeling remain unclear. Here, we investigated the role of the junctional kinase PAK4 in vertex remodeling. PAK4 localized to apical junctions and accumulated at multicellular vertices in MCDK cells and *Xenopus* embryos. Inhibition or knockout of PAK4 increased higher-order vertices, caused junctional discontinuities, and impaired barrier function in MDCK cells. PAK4 recruitment required the scaffolding protein Afadin. *Afdn*-KO cells exhibited severe junctional defects and reduced barrier function. Artificial targeting of PAK4 in *Afdn*-KO cells partially restored junctional continuity and barrier function. In *Xenopus* embryos, live imaging showed dynamic PAK4 accumulation at remodeling vertices, and PAK4 inhibition hindered resolution of multicellular vertices. Expression of an amino-terminal fragment (PAK4-NT) impaired remodeling and induced cytokinetic failure. Live imaging in *Xenopus* revealed barrier leakage at multicellular vertices upon PAK4 inhibition. These findings indicate that PAK4 and Afadin cooperate to maintain epithelial integrity and barrier function by promoting vertex remodeling.

**Summary:** Adhikary et al. show that PAK4, recruited to epithelial junctions by Afadin, drives dynamic vertex remodeling in cells and embryos. Inhibiting PAK4 disrupts tissue packing and compromises junctional continuity and barrier function, implicating PAK4 and Afadin as critical partners in vertex remodeling to preserve tissue integrity.

## Introduction

Epithelial tissues maintain their architecture and contribute to the homeostasis of multicellular organisms by forming well-ordered sheets of tightly connected epithelial cells. Dynamic rearrangement of epithelial sheets is essential not only during morphogenesis processes in development but also for the maintenance of adult tissues (Guillot and Lecuit, 2013b; Heisenberg and Bellaïche, 2013). In many epithelia, constituent cells are continuously turned over – old cells are removed from the sheet, while new cells are added either through cell division within the sheet or by insertion from outside the sheet. To preserve epithelial integrity and barrier function, the tissue must accommodate cell rearrangements such as intercalation, positional exchange between neighboring cells, and displacement, while maintaining intercellular adhesion and overall junctional organization (Varadarajan et al., 2019; Paré and Zallen, 2020; Mira-Osuna and Le Borgne, 2024; Higashi et al., 2024). Disruption of these processes compromises epithelial barrier integrity, which is critical for preventing uncontrolled paracellular flux and protecting against inflammation and pathogen invasion (Krug et al., 2014; Luissint et al., 2016; Saha et al., 2025).

Cell-cell junctions, composed of tight junctions (TJs), adherens junctions (AJs), and desmosomes, mediate intercellular adhesion in epithelial cells (Farquhar and Palade, 1963). TJs are crucial for maintaining the epithelial barrier (Van Itallie and Anderson, 2014; Otani and Furuse, 2020; Balda and Matter, 2023; Citi et al., 2024), whereas AJs and desmosomes provide mechanical strength (Meng and Takeichi, 2009; Rübsam et al., 2018; Eckert et al., 2025). To accommodate frequent cellular rearrangements, these junctional complexes undergo remodeling in response to epithelial reorganization, cell signaling, and cytoskeletal rearrangements, thereby supporting both tissue renewal and morphogenesis while preserving adhesion. Of particular interest is the remodeling that occurs at cell vertices – the points where multiple cell-cell junctions converge. In this study, we use the term “vertex” to describe such sites and specify them according to the number of participating cells: tricellular vertex (three cells), tetracellular or 4-way vertex (four cells), and rosette (five or more cells). We collectively refer to 4-way vertices and rosettes as higher-order vertices.

At these multicellular vertices of vertebrate epithelial cells, specialized junctional structures are assembled. Among them, tricellular tight junctions (tTJs) are well characterized, with distinct molecular components such as angulin family proteins and tricellulin, and tTJs play a crucial role in sealing the paracellular space of vertices (Ikenouchi et al., 2005; Masuda et al., 2011; Higashi et al., 2013; Sugawara et al., 2021). By contrast, the existence of tricellular adherens junctions (tAJs) has been proposed (Yonemura, 2011) and their components and regulation have recently begun to be understood in *Drosophila* (Lye et al., 2014; Finegan et al., 2019; Letizia et al., 2019; Uechi and Kuranaga, 2019), but the precise molecular composition and architecture of vertebrate tAJs remains less well defined.

Multicellular vertices are dynamically remodeled during tissue rearrangement (Sugimura and Otani, 2024). Several different modes of vertex remodeling have been characterized. One is vertex sliding, in which a tricellular vertex shifts position along a bicellular junctions as it shrinks or elongates (Vanderleest et al., 2018). Another is the T1 transition (Weaire and Rivier, 1984; Bertet et al., 2004), a topological transition in which two neighboring tricellular vertices converge as the intervening bicellular junction shortens, transiently forming a 4-way vertex. This intermediate state then resolves by elongation of a new junction in the perpendicular direction, giving rise to two new tricellular vertices with exchanged neighbors. A third example occurs during the final stages of cytokinesis in epithelial cells. As the cleavage furrow ingresses, a transient 4-way vertex is formed at the division plane. This intermediate state resolves either by establishing stable adhesion between the two daughter cells or by allowing the neighboring cells to insert between the daughters, thereby generating a new neighbor-neighbor interface (Gibson et al., 2006; Herszterg et al., 2013; Guillot and Lecuit, 2013a; Founounou et al., 2013; Higashi et al., 2016; Firmino et al., 2016; Landino et al., 2025). Such cytokinesis-associated remodeling closely resembles the later phase of a T1 transition and can thus be regarded as a specialized mode of vertex remodeling in proliferating epithelia. In addition, more complex rearrangements involve the formation of rosette structures, in which five or more cells converge at single vertex before being resolved into simpler configurations (Blankenship et al., 2006). These events are coordinated by adhesion complexes and the actomyosin cytoskeleton, under the control of Rho-family GTPases and their effectors (Lecuit and Yap, 2015; Arnold et al., 2017; Perez-Vale and Peifer, 2020; Huebner et al., 2021; Cavanaugh et al., 2022; Campàs et al., 2024). However, the molecular mechanisms that ensure barrier integrity during these highly dynamic vertex remodeling processes remain incompletely understood.

PAK4 is a serine/threonine kinase originally identified as a Cdc42 effector (Abo et al., 1998) and is essential for embryonic viability in mice (Qu et al., 2003). PAK4 has been implicated in apical junction formation (Wallace et al., 2010) and cell polarity (Selamat et al., 2015). Although PAK4 is localized at cell-cell junctions (Hou et al., 2013; Selamat et al., 2015) and its preferential localization at multicellular vertices has been observed (Baskaran et al., 2021), its roles in regulating junctional remodeling are not fully understood. Recently, proximity labeling studies identified Afadin, a scaffold protein essential for junctional actin organization (Mandai et al., 1997; Ikeda et al., 1999; Sakakibara et al., 2018, 2020; Kuno et al., 2025; Gong et al., 2025), as a possible binding partner of PAK4 (Baskaran et al., 2021; Choi et al., 2025). Afadin is required for the localization of PAK4 at cell-cell junctions (Baskaran et al., 2021) and for the maintenance of epithelial integrity (Choi et al., 2016; Sakakibara et al., 2020). Furthermore, the *Drosophila* homolog of Afadin, Canoe (Cno), preferentially localizes to epithelial vertices and has been linked to their dynamic remodeling events (Sawyer et al., 2009; Choi et al., 2013; Yu and Zallen, 2020), suggesting that PAK4 may contribute to the regulation of vertex remodeling in coordination with Afadin.

In this study, we investigated the function and dynamics of PAK4 at epithelial vertices using gene-edited MDCK II cells and live imaging of *Xenopus laevis* embryos. We further examined the role of Afadin in recruiting PAK4 to junctions and demonstrate that both proteins are critical for maintaining epithelial integrity and barrier function specifically at cell vertices.

## Results

### PAK4 preferentially accumulates at higher-order vertices

To examine the localization of PAK4 in epithelia that undergo dynamic remodeling *in vivo*, frozen sections of mouse small intestine were stained with an anti-PAK4 antibody. PAK4 was detected at cell-cell junctions and was particularly enriched at 4-way (tetracellular) vertices and at some tricellular vertices (Fig. 1A). Quantitative analysis revealed that PAK4 accumulation at tricellular vertices was increased compared to the average intensity at bicellular junctions (1.45 ± 0.65-fold) (Fig. 1B). At 4-way vertices, PAK4 was enriched to approximately twice the level observed at bicellular junctions (2.23 ± 1.13-fold) (Fig. 1B). The TJ-associated protein ZO-1 was also enriched at tricellular vertices (1.48 ± 0.65-fold) and 4-way vertices (1.83 ± 0.54-fold) (Fig. 1C), although its accumulation at 4-way vertices was generally less prominent than that of PAK4 (Fig. 1A, right panels).

**Figure 1.**
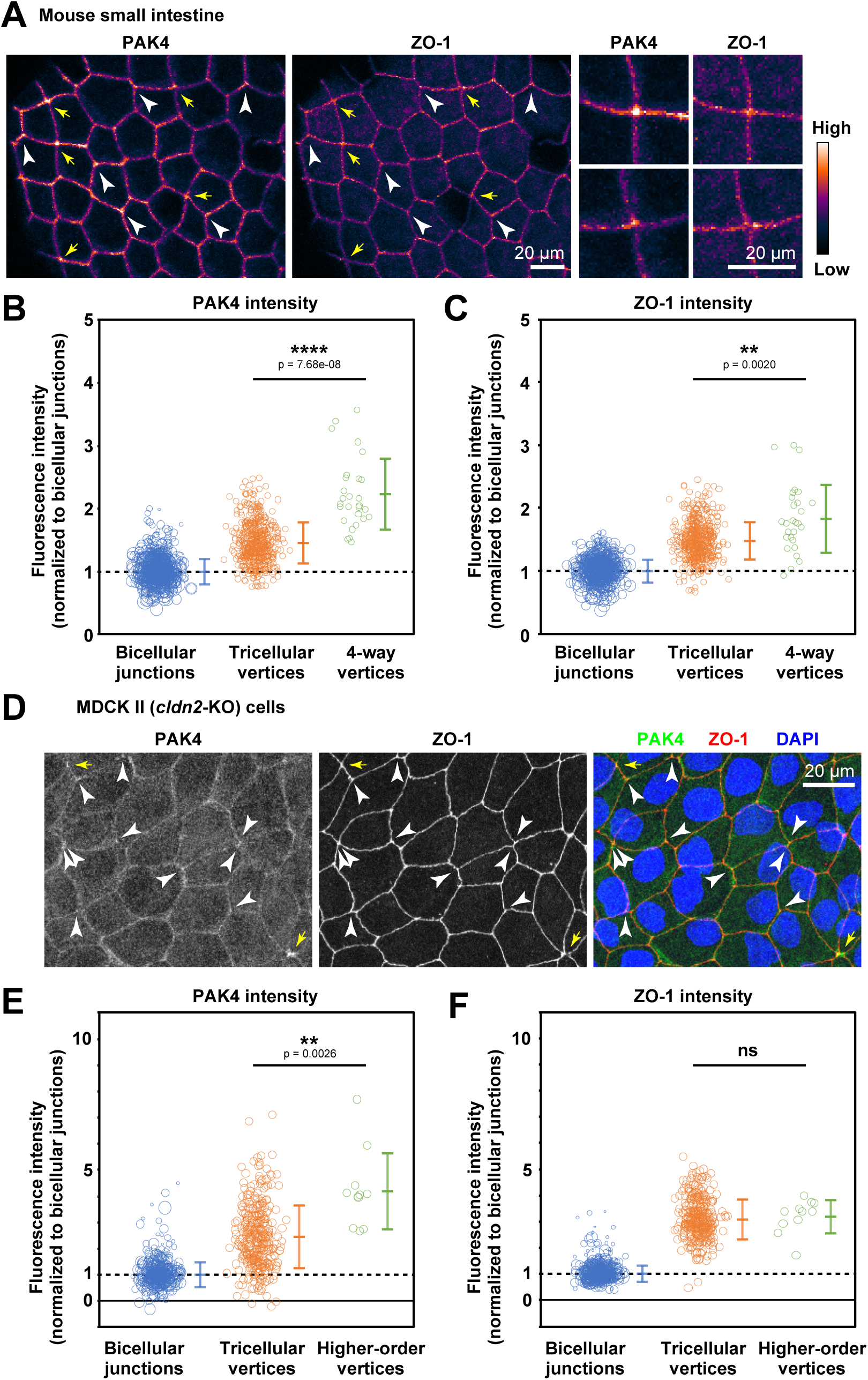
PAK4 is enriched at higher-order cell-cell junctions in the mouse small intestine and MDCK II cells. (A,D) Immunofluorescence staining of mouse small intestine fixed with formaldehyde (A) and of EtOH-fixed Ctrl MDCK II cells (D). Samples were labeled with rabbit anti-PAK4 mAb and mouse anti-ZO-1 mAb (A) or rabbit anti-PAK4 mAb (green), rat anti-ZO-1 mAb (red) and DAPI (blue) (D). White arrowhead, tricellular vertices with PAK4 signal; yellow arrows, 4-way vertices/rosettes. Scale bars, 20 µm. (B, C, E, F) Quantification of fluorescence intensities. Background-subtracted fluorescence intensities of PAK4 (B, E) and ZO-1 (C, F) at tricellular vertices and 4-way vertices/rosettes were normalized to the junction-length-weighted average intensity at bicellular junctions. Individual measurements at bicellular junctions are shown as circles, with the areas proportional to junction length. Individual measurements at tricellular vertices and 4-way vertices/rosettes are shown as circles and the bars indicate mean ± sd. The values of tricellular vertices and 4-way vertices/rosettes were compared using a two-tailed unpaired Welch’s *t*-test (ns, p > 0.05; **, p < 0.01; ****, p < 0.0001).

In cultured *cldn2*-KO MDCK II (Ctrl) cells (Saito et al., 2021), PAK4 was localized at bicellular junctions, as previously reported (Hou et al., 2013; Selamat et al., 2015; Baskaran et al., 2021) and showed enrichment at 4-way vertices/rosettes and many tricellular vertices (Fig. 1D). Quantification of fluorescence intensity revealed that PAK4 accumulated at tricellular vertices to varying degrees, in most cases ranging from levels similar to those at bicellular junctions to as much as fivefold higher (2.45 ± 1.20-fold) (Fig. 1E). Additionally, some tricellular vertices showed marked enrichment (Fig. 1D, arrowheads; Fig. 1E). Moreover, at 4-way vertices and rosettes (Fig. 1D, arrows), PAK4 was consistently enriched on average to approximately four times the level observed at bicellular junctions (4.22 ± 1.48-fold) (Fig. 1E). By comparison, ZO-1 showed relatively uniform accumulation, reaching about threefold the level of bicellular junctions at both tricellular (3.11 ± 0.76-fold) and 4-way vertices/rosettes (3.21 ± 0.64-fold) (Fig. 1F).

These results indicate that PAK4 is enriched at multicellular vertices of cell-cell junctions in mammalian epithelial cells.

### PAK4 accumulates at remodeling vertices in Xenopus gastrula-stage epithelia

To examine the dynamics of PAK4 at remodeling vertices in a developing organism, we turned to gastrula-stage *Xenopus laevis* embryos. Immunostaining with an anti-PAK4 antibody detected PAK4 at cell-cell junctions, and PAK4 was particularly enriched at higher-order vertices (4-way vertices and rosettes) as well as at some tricellular vertices (Fig. 2A). We then live imaged PAK4-mNeon with TJ (TagBFP-ZO-1)- and AJ (PLEKHA7-mCherry)-localizing proteins (Fig. 2B). Quantitative analysis of live images revealed that PAK4 accumulation at 4-way vertices and rosettes (2.74 ± 0.21) was significantly higher than at tricellular vertices (1.64 ± 0.11) (Fig. 2C). ZO-1 was also enriched at 4-way vertices and rosettes (3.47 ± 0.25) compared to tricellular vertices (1.84 ± 0.13) (Fig. 2D). To examine the lateral localization of PAK4, we quantified the relative fluorescence intensity along the *z*-axis (Fig. 2E) (Higashi et al., 2019). The PAK4-mNeon peak closely overlapped with the ZO-1 peak, while it was slightly offset from the PLEKHA7 peak, indicating that PAK4 co-localizes with apical junctional proteins (Fig. 2E).

**Figure 2.**
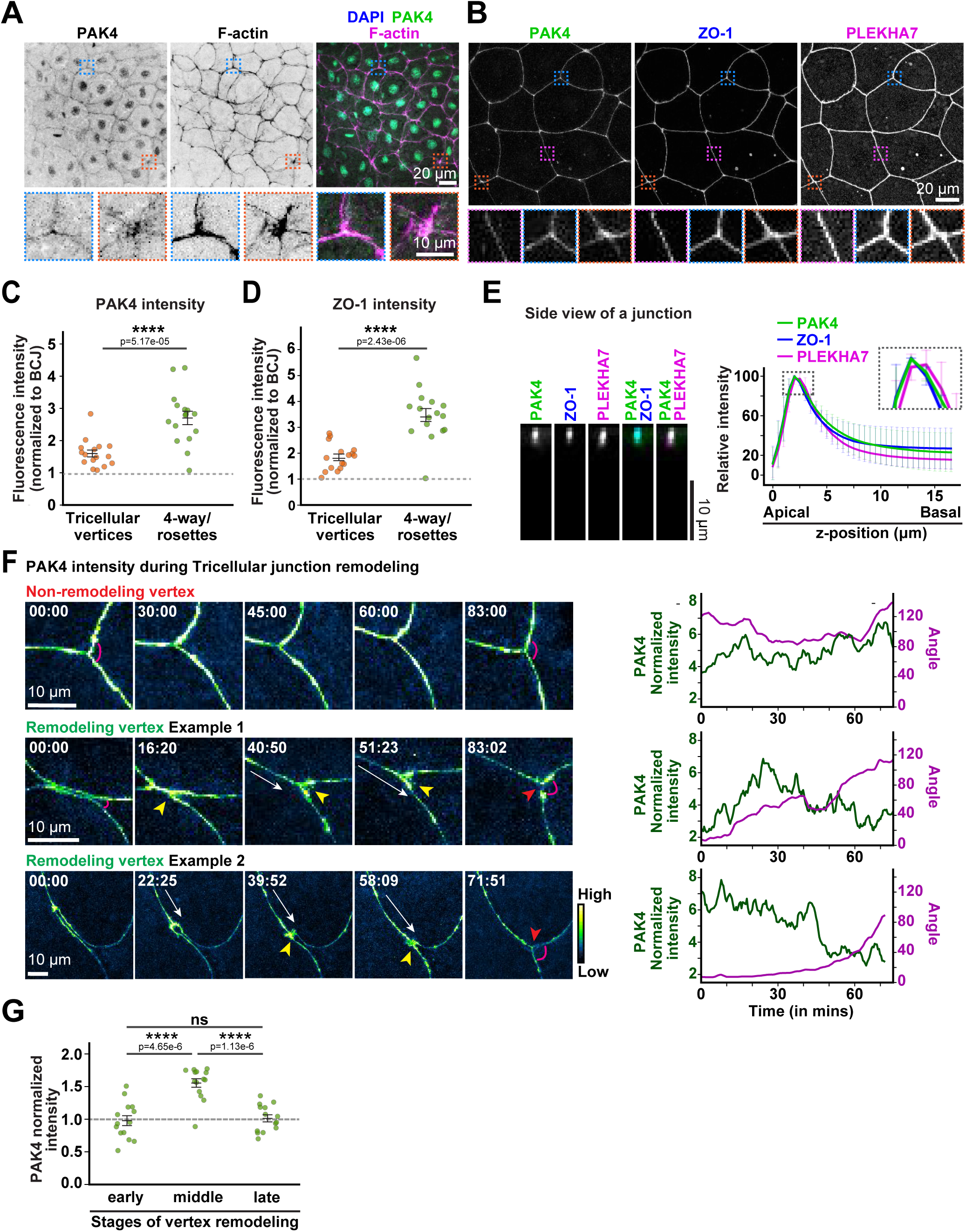
PAK4 is enriched at remodeling higher-order cell-cell vertices in *Xenopus* epithelia. (A) Immunofluorescence staining of the animal hemisphere of gastrula-stage (NF stage 10) *Xenopus* embryos with mouse anti-PAK4 mAb (green), phalloidin (magenta), and DAPI (blue). Zoomed-in panels highlight PAK4 recruitment at tricellular vertices (blue boxes) and 4-way vertex/rosette (orange boxes). Scale bars, 20 µm (upper), 10 µm (lower). (B) Live confocal images of epithelial cells in gastrula-stage *Xenopus* embryo expressing mNeon-PAK4 (PAK4), TagBFP2-ZO-1 (ZO-1), and PLEKHA7-mCherry (PLEKHA7). Zoomed-in panels highlight localization of PAK4, ZO-1, and PLEKHA7 at bicellular junction (magenta), tricellular vertex (blue), and higher-order vertex (orange). Scale bar, 20 µm. (C,D) Quantification of PAK4 (C) and ZO-1 (D) fluorescence intensities at tricellular vertices and 4-way vertices/rosettes, normalized to bicellular junction (BCJ) fluorescence. Individual measurements (circles) and mean ± SEM (bars) are shown. Values were compared using a two-tailed unpaired Student’s *t*-test (****, p < 0.0001; n = 16 vertices, 4 embryos). (E) Representative side view of a bicellular junction. PAK4 (green) is localized apically along with TJ protein ZO-1 (blue). It was a slight offset from AJ protein PLEKHA7 (magenta). Scale bar, 10 µm. Graph shows intensity profiles of PAK4, ZO-1, and PLEKHA7 along the apico-basal axis (mean ± SD. n = 16 junctions, 5 embryos). (F) Montage of non-remodeling (top) and remodeling (middle and bottom) vertices labeled with PAK4-mNeon, shown using the green Fire blue lookup table (LUT). PAK4 intensity at the vertex (green line on graphs) and vertex angle (magenta line on graphs) were measured over time. White arrows, vertex movement; yellow arrowheads, PAK4 accumulation; red arrowheads, end of remodeling. Scale bars, 10 µm. (G) Quantification of PAK4 intensity at remodeling vertices at early, middle, and late stages of remodeling, normalized to the average intensity of non-remodeling vertices at the same time points. Individual measurements (circles) and mean ± SEM (bars) are shown. Values were compared using a two-tailed unpaired Student’s *t*-test (****, p < 0.0001; n = 14 vertices, 7 embryos, 3 independent experiments).

To understand dynamic changes in PAK4 localization during junction remodeling, we performed live imaging of PAK4-mNeon at remodeling and non-remodeling vertices in the gastrula-stage *Xenopus* epithelium. We defined a “remodeling vertex” as a tricellular vertex that has an acute angle to start off, and then actively remodels over several minutes, resulting in a vertex angle greater than 90°. In contrast, we defined a “non-remodeling vertex” as a tricellular vertex where the starting angle is >90°, and the change in angle is ± 20°. PAK4 appeared to be enriched at actively remodeling vertices, whereas PAK4 intensity remained more stable at non-remodeling vertices (Fig. 2F and Video 1). Indeed, quantification showed that early in vertex remodeling, PAK4 intensity was similar at remodeling vs. non-remodeling vertices. Then, midway through vertex remodeling PAK4 at remodeling vertices was significantly enriched (1.56 ± 0.07-fold) over PAK4 at non-remodeling vertices (Fig. 2G). PAK4 intensity at remodeling vertices then returned to baseline levels as remodeling completed (Fig. 2G).

These results indicate that PAK4 is enriched at multicellular vertices in developing epithelial tissue, and PAK4 especially accumulates at remodeling tricellular vertices.

### Inhibition of PAK4 disrupts vertex geometry in MDCK II epithelial sheets

To assess the function of PAK4, Ctrl MDCK II cells were treated with different concentrations of the ATP-competitive PAK4 inhibitor, PF-3758309. Treatment with PAK4 inhibitor led to an increased number of higher-order vertices compared to DMSO-treated cells (Fig. 3A). We performed three types of quantitative analyses to evaluate this effect. First, the ratio of the total number of cell vertices to the total number of cells in the monolayer was calculated. In an ideal hexagonally packed epithelial sheet, each vertex is formed where three cells meet, and each cell contributes to six such vertices. Given this geometry, the total number of vertices divided by the total number of cells is expected to be 2. When cells deviate from hexagonal packing and higher-order junctions (involving four or more cells) are present, this ratio decreases. For example, in tightly packed arrays of rectangular and triangular cells, the vertex-to-cell ratios become 1 and 0.5, respectively. Discontinuities in cell-cell junctions can also lead to a further reduction in the number of detectable vertices. In this analysis, monolayers treated with DMSO or 500 nM PAK4 inhibitor exhibited vertex-to-cell ratios close to 2 (1.82 ± 0.05 [DMSO] and 1.80 ± 0.09 [500 nM PAK4 inhibitor]) (Fig. 3B). In contrast, treatment with 5 µM PAK4 inhibitor resulted in a significant reduction in this ratio (1.71 ± 0.08) (Fig. 3B), suggesting a disturbance in normal junctional organization. Second, all vertices were classified according to the number of converging cells, and the proportions of 4-way vertices and rosettes were quantified. At 500 nM PAK4 inhibitor, a significant increase in 4-way vertices was observed, and at 5 µM, both 4-way vertices and rosettes were significantly increased (Fig. 3C). Third, the number of discontinuities in cell-cell junctions was quantified. A slight increase was noted in the 5 µM PAK4 inhibitor-treated cells, although the change was not statistically significant (Fig. 3D).

**Figure 3.**
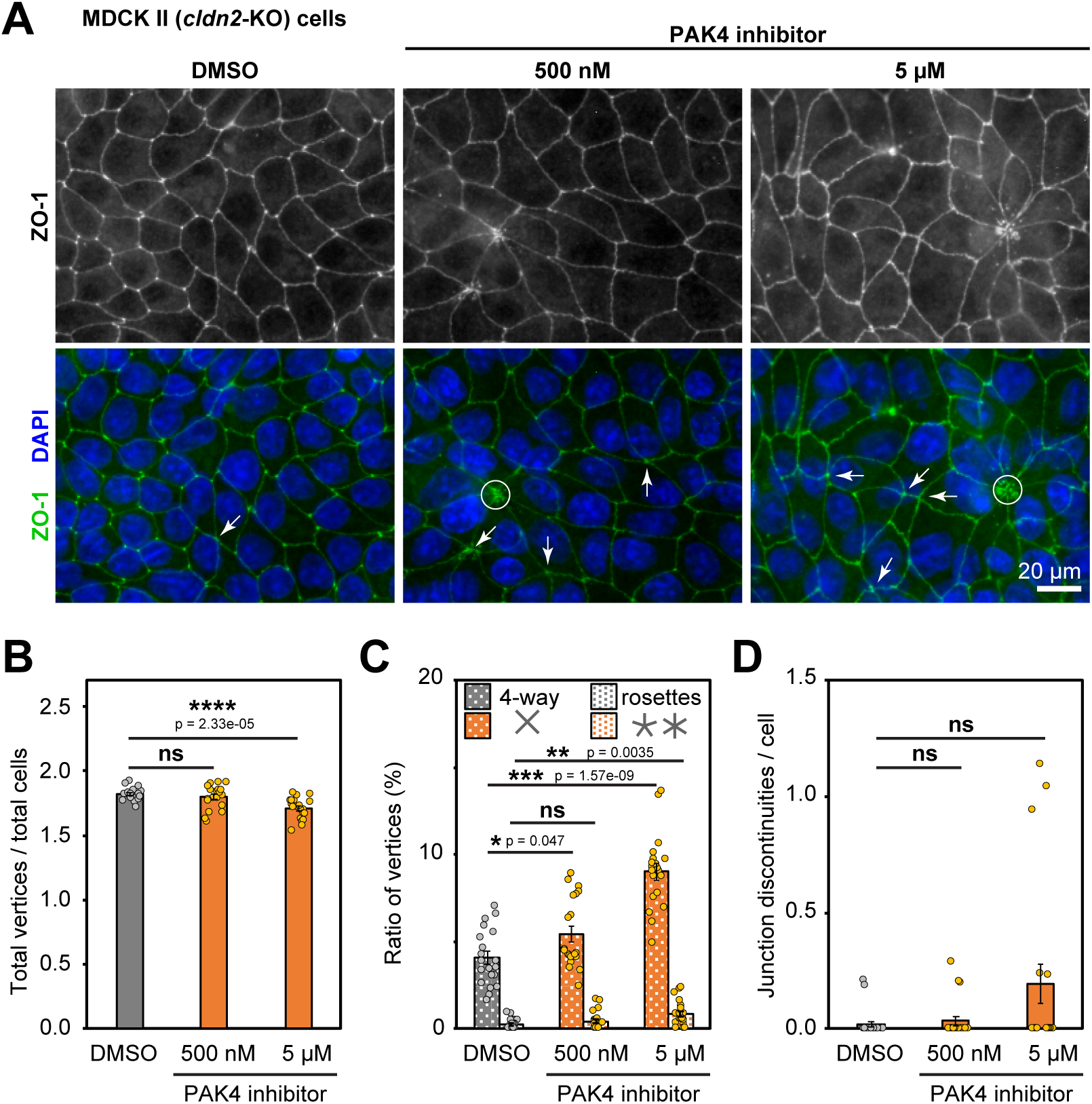
PAK4 inhibition leads to an increase in higher-order cell-cell vertices. (A) Immunofluorescence staining of DMSO- or PF-3758309 (PAK4 inhibitor)-treated Ctrl MDCK II (*cldn2*-KO) cells with rat anti-ZO-1 mAb (green) and DAPI (blue). Arrows, 4-way junctions; circle, rosettes. Scale bar, 20 µm. (B-D) Quantification of vertex-to-cell ratio (B), proportions of 4-way vertices and rosettes (C), and number of discontinuities in cell-cell junctions (D). The values in PAK4 inhibitor-treated monolayers (n = 20) were compared with those of DMSO-treated monolayers (n = 20) using a two-tailed unpaired Welch’s *t* test with Bonferroni’s correction (ns, p > 0.05; *, p < 0.05; **, p < 0.01; ***, p < 0.001; p < 0.0001). S.E.M. (bars) and individual measurements (circles) are shown.

These results suggest that kinase activity of PAK4 is required for the maintenance of tricellular vertices as well as for the resolution of 4-way vertices and rosettes.

### PAK4 inhibition hinders resolution of higher-order vertices in *Xenopus* epithelium

To assess the function of PAK4 during vertex remodeling in development, *Xenopus* embryos expressing TagBFP2-ZO-1, Angulin-1-3xGFP, and the F-actin probe LifeAct-miRFP were treated with the PAK4 inhibitor, PF-3758309. PAK4 inhibition led to abnormal clumpy signal around multicellular vertices for ZO-1, Angulin-1, and F-actin compared with control DMSO-treated embryos (Fig. 4A). To investigate the resolution of multicellular vertices in developing embryos, we examined tissue packing following cytokinesis. Daughter cells in control (DMSO-treated) and PAK4 inhibitor-treated embryos were categorized based on the geometry of the newly formed interface: daughter-daughter interface (Type I), 4-way vertex (Type II), or neighbor-neighbor interface (Type III) (Fig. 4B) (Gibson et al., 2006; Higashi et al., 2016). Control embryos displayed a 4-way vertex (Type II, 100%) immediately after cytokinesis, and after 60 minutes, some of these rearranged into daughter-daughter interfaces (Type I, 68.4%), while others rearranged into neighbor-neighbor interfaces (Type III, 10.5%) (Fig. 4C-D). PAK4 inhibitor-treated embryos also showed a high percentage of 4-way vertices (Type II, 95.7%) immediately after cytokinesis, but the majority of these failed to rearrange over time; 69.6% remained as 4-way vertices 60 minutes post cytokinesis, while 13.0% rearranged into daughter-daughter interfaces (Type I), and 17.4% rearranged into neighbor-neighbor interfaces (Type III) (Fig. 4C-D). Therefore, PAK4 inhibition hinders 4-way vertex resolution following cytokinesis, affecting cell packing patterns in the developing *Xenopus* epithelium.

**Figure 4.**
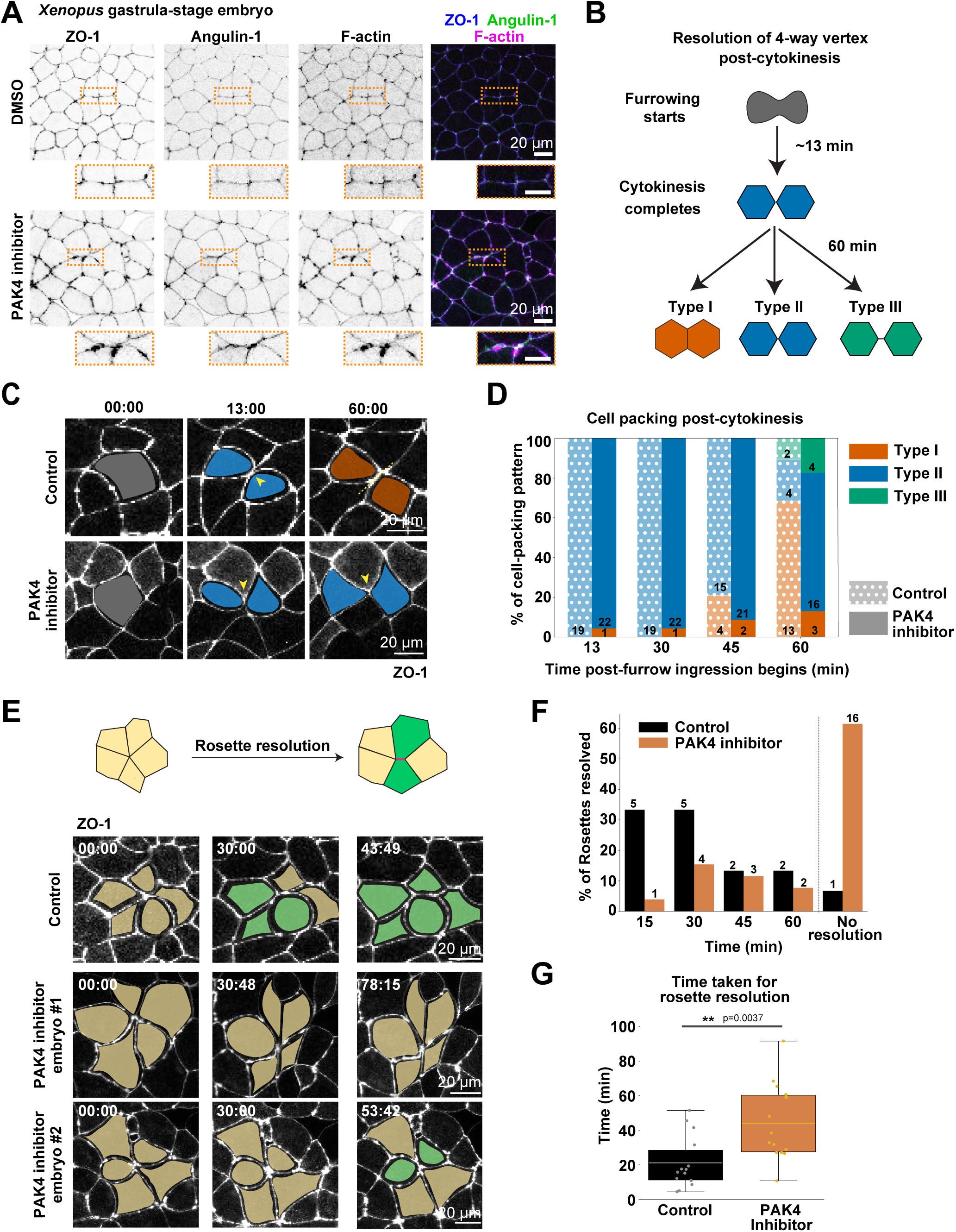
PAK4 inhibition impairs higher-order junction remodeling in *Xenopus* epithelia. (A) Live confocal images of epithelial cells in the animal hemisphere of gastrula-stage *Xenopus* embryos expressing TagBFP2-ZO-1 (ZO-1), Angulin-1-3xGFP (Angulin-1), and LifeAct-miRFP (F-actin) treated with 0.1% DMSO (control) or 10 µM PAK4 inhibitor PF-3758309. Zoomed-in panels (orange boxes) highlight the junctional localization pattern of ZO-1, Angulin-1, and F-actin in control versus PAK4 inhibitor-treated cells. Scale bars, 20 µm. (B) Schematic of 4-way vertex resolution following cytokinesis. (C) Montage showing cells before cleavage furrow ingression (gray) and at 13 and 60 min after ingression. Dividing cells are color coded for type I (orange) and type II (blue) interfaces. Scale bars, 20 µm. (D) Quantification of types I, II, and III interfaces at 13, 30, 45, and 60 min after ingression. (control: 3 embryos; inhibitor: 4 embryos; 3 independent experiments). (E) Montage of control and PAK4 inhibitor-treated embryos expressing TagBFP2-ZO-1, showing rosettes before (yellow) and after resolution (green). Scale bars, 20 µm. (F) Quantification of rosette resolution at 15, 30, 45, and 60 min after the onset of imaging (control: 5 embryos, 15 rosettes; inhibitor: 6 embryos, 26 rosettes; 3 independent experiments). (G) Quantification of time required for rosette resolution (control: 5 embryos, 14 rosettes; inhibitor: 6 embryos, 14 rosettes; 3 independent experiments). Data were compared using a two-tailed unpaired Student’s *t*-test (**, p < 0.01).

Next, we examined whether PAK4 inhibition causes defects in rosette resolution in developing *Xenopus* epithelia. Rosette resolution was defined as the rearrangement of a multicellular rosette – where five or more cells meet – into a simpler configuration, in which newly adjoining neighbors extend their bicellular junctions and generate tricellular or 4-way vertices. PAK4 inhibition causes a significant delay in rosette resolution compared to DMSO-treated control embryos (Fig. 4E and Video 2). In control embryos, the majority of rosettes resolve within 30 minutes (66.67%), with only a small percentage (6.67%) not resolved after 60 minutes of imaging (Fig. 4F). In contrast, in PAK4 inhibitor-treated embryos, the majority of rosettes (61.54%) were not resolved after 60 minutes of observation (Fig. 4F). In the population of rosettes that were resolved, the average time for resolution was significantly higher in PAK4 inhibitor-treated embryos (median = 35.83 min) compared to controls (median = 17.13 min) (Fig. 4G). A limitation of this rosette resolution analysis is that we quantified rosettes from the start of imaging rather than from the time of their actual formation, which leads to an underestimation of resolution times. However, this systematic bias was applied equally to both control and PAK4 inhibitor-treated embryos.

Taken together, these results suggest that PAK4 kinase activity is required for proper organization of junctional proteins and F-actin at multicellular vertices and plays a crucial role in resolution of 4-way vertices and rosettes in developing *Xenopus* epithelia.

### PAK4-knockout epithelial sheets exhibit discontinuities in cell-cell junctions and an increased number of multicellular vertices

To further examine the function of PAK4, we established two independent *PAK4*-knockout (KO) MDCK II cell clones (*PAK4*-KO#1 and *PAK4*-KO#2) with CRISPR/Cas9-based genome editing (Fig. S1A-C) (Saito et al., 2024) using Ctrl cells as a parental cell line. Immunostaining and immunoblotting analyses confirmed complete loss of PAK4 signal in both clones (Fig. S1G-H).

Using these *PAK4*-KO cell clones, we examined the roles of PAK4 at cell-cell junctions and vertices. When cultured on a glass surface, Ctrl cells formed a well-ordered monolayer by 40 hours after plating. In contrast, *PAK4*-KO cells formed disorganized sheets, with both clones showing a significant increase in the number of discontinuities in ZO-1 signal (Fig. 5A). In *PAK4*-KO cells, the vertex-to-cell ratio was significantly decreased to less than 1.0 (Fig. 5B), and the proportion of 4-way vertices and rosettes among all cell vertices was increased compared to Ctrl cells, although the increase reached statistical significance only in one clone (*PAK4*-KO#1) (Fig. 5C). Additionally, the number of junction discontinuities was around 0.5 per cell, which was much higher than the Ctrl (Fig. 5D). Stable expression of Flag-tagged wild-type PAK4 in the *PAK4*-KO#1 clone restored both vertex organization and junctional continuity (Fig. S2A-E). In contrast, expression of a kinase-dead mutant (Flag-PAK4 K351R), which is equivalent to a K299R mutation in PAK1 (Zhang et al., 1995; Sells et al., 1997), failed to rescue the discontinuity phenotype (Fig. S2A-E), suggesting that kinase activity is required for maintaining epithelial integrity. Vertex organization was only partially restored by Flag-PAK4 K351R, implicating a kinase activity-independent contribution of PAK4 to vertex patterning.

**Figure 5.**
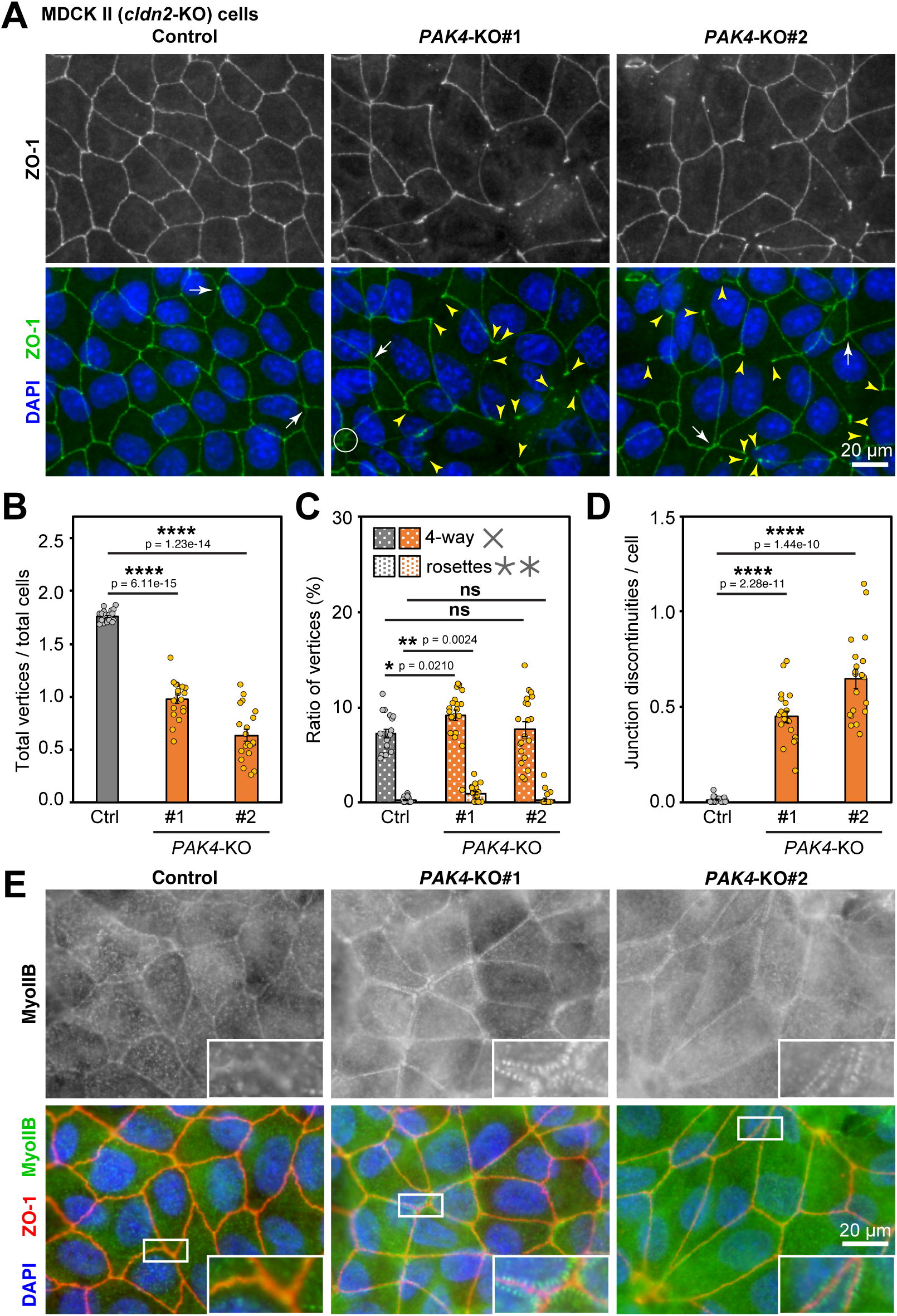
Loss of PAK4 impairs epithelial integrity in cultured cells. (A) Immunofluorescence staining of Ctrl and *PAK4*-KO cell clones (#1 and #2) with anti-ZO-1 mAb (green) and DAPI (blue). Arrows, 4-way vertices; circle, rosettes; yellow arrowheads, discontinuities in cell-cell junctions. Scale bar, 20 µm. (B-D) Quantification of the vertex-to-cell ratio (B), proportions of 4-way vertices and rosettes (C), and number of discontinuities in cell-cell junctions (D). Values from *PAK4*-KO monolayers (n = 20) were compared with those of Ctrl monolayers (n = 20) using a two-tailed Welch’s *t*-test with Bonferroni’s correction (ns, p > 0.05; *, p < 0.05; **, p < 0.01; ****, p < 0.0001). S.E.M. (bars) and individual measurements (circles) are shown. (E) Immunofluorescence staining of Ctrl and *PAK4*-KO cells with anti-MyoIIB pAb (green), anti-ZO-1 mAb (red), and DAPI (blue). Scale bar, 20 µm.

These results with *PAK4*-KO cells complement our experiments with the PAK4 inhibitor, further supporting the role of PAK4 in maintaining junctional integrity in epithelial cells.

### PAK4 knockout results in formation of sarcomere-like MyoIIB structures along cell-cell junctions

We noticed that the cell-cell junctions of *PAK4*-KO MDCK II cells cultured on glass coverslips appear straighter than Ctrl cells, and this effect was more obvious when the cells were cultured on polycarbonate filters (Fig. S3D). Since the shape of cell-cell junctions is determined by the actomyosin cytoskeleton, we next examined the Myosin II organization in *PAK4*-KO cells. Whereas Ctrl cells exhibit faint staining for MyoIIB along cell-cell junctions, *PAK4*-KO cells show dotted MyoIIB signal flanking both sides of cell-cell junctions (Fig. 5E), which closely resembles the sarcomere-like structure reported in inner ear cells (Ebrahim et al., 2013), ZO-1/2-double-deficient MDCK II cells (Yamazaki et al., 2008; Fanning et al., 2012; Choi et al., 2016), and *claudin/JAM-A*-KO MDCK II cells (Nguyen et al., 2024). MyoIIA also exhibited a punctate pattern at both sides of *PAK4*-KO cell-cell junctions (Fig. S3A) similar to MyoIIB (Fig. S3B). To examine whether junctional Myosin is activated, we stained for S18/T19-double phosphorylated myosin light chain (2P-MLC). 2P-MLC was localized at cell-cell junctions around the tricellular vertices in *PAK4*-KO cells (Fig. S3C), suggesting that the junctional tension is increased. Vinculin, an actin-binding protein recruited to AJs under tension (Yonemura et al., 2010; le Duc et al., 2010; Huang et al., 2017; Higashi et al., 2016; van den Goor et al., 2024), also accumulated at the tricellular region of *PAK4*-KO cells (Fig. S3D), while other TJ and AJ proteins did not exhibit notable changes in localization at junctions in *PAK4*-KO cells (Fig. S3E-H).

These results suggest that PAK4 suppresses the formation of a high tension actomyosin array flanking cell-cell junctions, thus regulating the linearity of cell-cell junctions.

### Afadin is required for the junctional localization of PAK4 and maintenance of epithelial integrity

It was reported that Afadin is responsible for localization of PAK4 at cell-cell junctions (Baskaran et al., 2021). To assess Afadin’s role in PAK4-driven vertex remodeling, we established Afadin knockout cell clones using CRISPR/Cas9 genome editing (*Afdn*-KO#1 and *Afdn*-KO#2) (Fig S1D-H) using Ctrl cells as a parental cell line. PAK4 was no longer localized at cell-cell junctions in *Afdn*-KO cells (Fig. S1G), consistent with the previous report (Baskaran et al., 2021). However, Afadin was still localized at cell-cell junctions and cell vertices of *PAK4*-KO cells (Fig. S1G), suggesting that Afadin recruits PAK4 to cell-cell junctions. It was reported that knockdown of Afadin in ZO-1/ZO-2-double knockdown MDCK II cells resulted in discontinuities at cell-cell junctions (Choi et al., 2016). To assess whether Afadin (independent of ZO-1/ZO-2 double knockdown) is required for maintenance of epithelial integrity, *Afdn*-KO cells were cultured on glass coverslips. Similar to *PAK4*-KO cells, both clones of *Afdn*-KO cells exhibited discontinuities in cell-cell junctions and an increased frequency of 4-way junctions (Fig. 6A-D). These phenotypes were largely rescued by exogenous expression of FLAG-Afadin in the *Afdn*-KO#1 cell clone (Fig. S2F-J).

**Figure 6.**
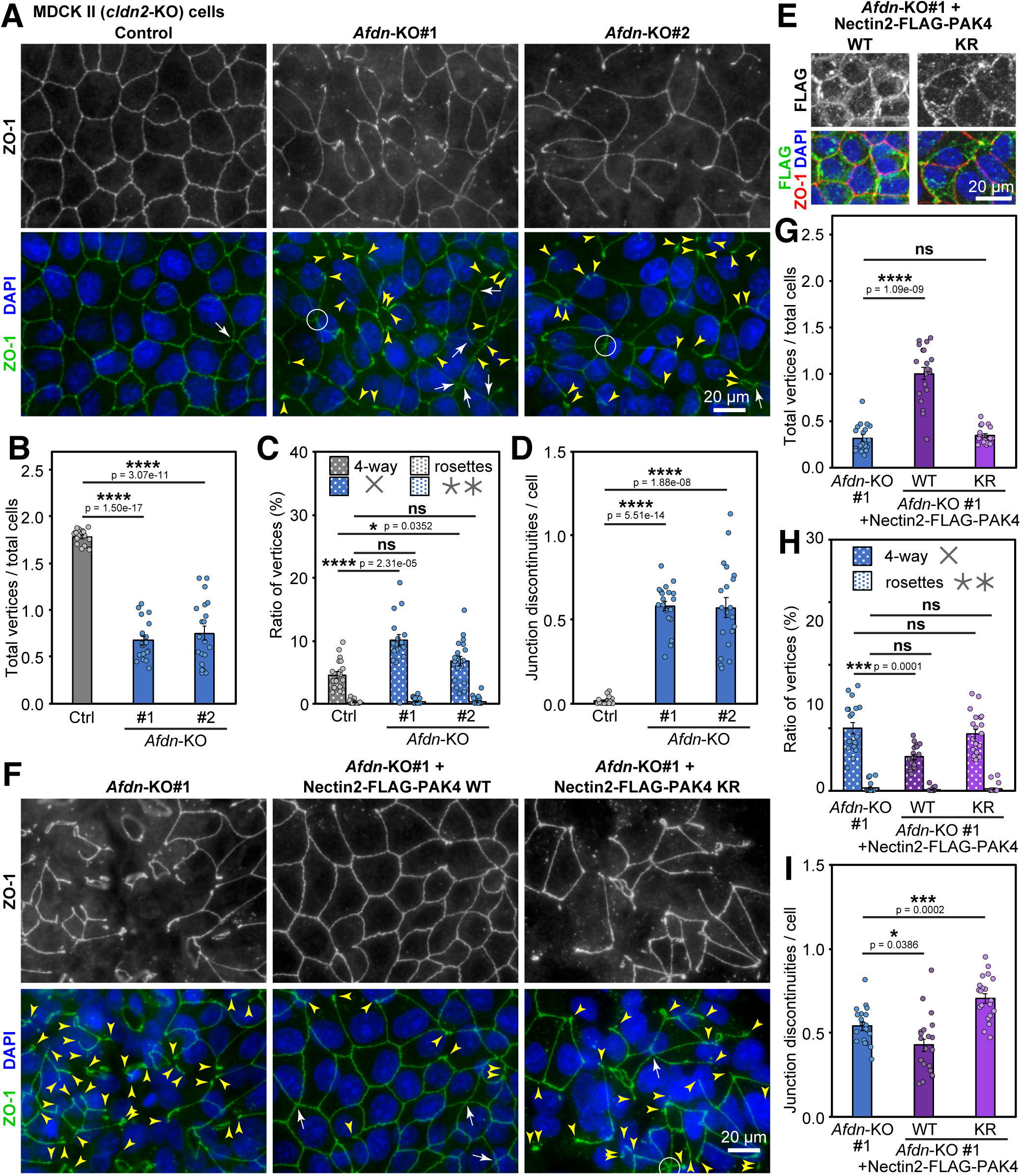
Loss of Afadin compromises epithelial integrity, partially rescued by junctional targeting of PAK4. (A) Immunofluorescence staining of Ctrl and *Afdn*-KO cell clones (#1 and #2) using anti-ZO-1 mAb (green) and DAPI (blue). Arrows, 4-way vertices; circle, rosette; yellow arrowheads, discontinuities in cell-cell junctions. Scale bar, 20 µm. (B-D) Quantification of the vertex-to-cell ratio (B), proportions of 4-way vertices and rosettes (C), and number of discontinuities in cell-cell junctions (D). Values from *Afdn*-KO monolayers (n = 20) were compared with those of Ctrl monolayers (n = 20) using a two-tailed Welch’s *t*-test with Bonferroni’s correction (ns, p > 0.05; *, p < 0.05; ****, p < 0.0001). S.E.M. (bars) and individual measurements (circles) are shown. (E) Immunofluorescence staining of *Afdn*-KO #1 cells expressing Nectin2-FLAG-PAK4 WT or KR mutant using anti-ZO-1 (red) and anti-FLAG (green) mAbs and DAPI (blue). Scale bar, 20 µm. (F) Immunofluorescence staining of *Afdn*-KO #1 cells and *Afdn*-KO #1 cells expressing Nectin2-FLAG-PAK4 WT or KR mutant labeled as in (A). Scale bar, 20 µm. (G-I) Quantification of the vertex-to-cell ratio (G), proportions of 4-way vertices and rosettes (H), and number of discontinuities in cell-cell junctions (I). Values from Nectin2-FLAG-PAK4-expressing Afdn-KO #1 cell monolayers (n = 20) were compared with those of *Afdn*-KO #1 monolayers (n = 20) using a two-tailed Welch’s *t*-test with Bonferroni’s correction (ns, p > 0.05; *, p < 0.05; ***, p < 0.001; ****, p < 0.0001). S.E.M. (bars) and individual measurements (circles) are shown.

These results suggest that Afadin is required for the recruitment of PAK4 to cell-cell junctions and the maintenance of epithelial integrity and vertex organization.

### Artificial targeting of PAK4 to adherens junctions partially rescues Afadin knockout phenotypes

Afadin is known for its role as an actin-binding protein that regulates the actin cytoskeleton at cell-cell junctions (Mandai et al., 1997). Additionally, we and others have shown that Afadin is required for PAK4 localization at cell-cell junctions (Fig. S1G) (Baskaran et al., 2021). Thus, we next examined how Afadin’s ability to recruit PAK4 to cell-cell junctions affects the maintenance of epithelial integrity. To test this, we assessed whether artificial recruitment of PAK4 to junctions is sufficient to maintain epithelial integrity in the absence of Afadin. We established an *Afdn*-KO cell clone stably expressing PAK4 conjugated with Nectin2, an Afadin-binding AJ adhesion protein (Takahashi et al., 1999). The Nectin2-FLAG-PAK4 construct was localized at cell-cell boundaries (Fig. 6E), but its localization was not restricted to AJs. Remarkably, the epithelial integrity defects of *Afdn*-KO cells were partially rescued in Nectin2-FLAG-PAK4-expressing cells (Fig. 6F-I). The vertex-to-cell ratio was significantly increased to around 1.0 (Fig. 6G), and the frequency of 4-way junctions was significantly decreased compared to the *Afdn*-KO cells (Fig. 6H). In addition, the frequency of junctional discontinuities was significantly reduced (Fig. 6I). Notably, the kinase-dead version of PAK4 conjugated with Nectin2 failed to rescue the phenotypes (Fig. 6F-I).

Together, these data suggest that junctional localization of PAK4 is important for the maintenance of epithelial integrity and that one of Afadin’s key roles is to recruit PAK4 to cell-cell junctions and cell vertices.

### Afadin interacts with the N-terminal region of PAK4

To gain insights into the binding interface between PAK4 and Afadin, we used AlphaFold3 (Abramson et al., 2024) to predict the 3D structure of the PAK4-Afadin complex (Fig. S4A). Although a large part of the PAK4 and Afadin polypeptides appeared unstructured, the amino-terminal region of PAK4 near the p21-binding domain (PBD) (PAK4-NT; aa 68-89) and the amino-terminal part of Afadin including the Ras association 1 (RA1) domain exhibited low predicted aligned error (PAE) scores (Fig. S4B), suggesting that these regions may interact directly. Further, the AlphaBridge algorithm (Álvarez-Salmoral et al., 2024) suggested possible interaction of these regions (Fig. S4C,F). The interaction of the predicted Afadin-interacting region of PAK4 with Afadin appears to affect the conformation of the adjacent PAK4 autoinhibitory domain (AID), which is associated with the active site groove of the kinase domain in the inactive state (Fig. S4B,D-E). Interestingly, the amino-acids involved in the interaction in the Afadin-PAK4 complex model are at least partially conserved among the group 2 PAK family including PAK4, PAK5, PAK6, and *Drosophila* mbt, but are not conserved in the group 1 PAK family including PAK1, PAK2, and PAK3 (Fig. S4G).

To examine whether the identified domains are responsible for interaction of PAK4 and Afadin, HEK293T cells were transfected with a GFP-tagged RA1 fragment of Afadin (GFP-Afdn-RA1) and mCherry-tagged full-length PAK4 (mChe-PAK4), and GFP-Afdn-RA1 was immunoprecipitated. A small amount of mChe-PAK4 was co-immunoprecipitated (Fig. S4H), suggesting that the Afadin RA1 domain can interact with PAK4. FLAG-tagged full-length Afadin could also be co-immunoprecipitated with GFP-tagged PAK4-NT (Fig. S4H).

These results implicate that PAK4 and Afadin may interact through the N-terminal region of PAK4 and the RA1 domain of Afadin.

### Inhibiting PAK4 recruitment to junctions prevents vertex remodeling and causes cytokinetic failure in the *Xenopus* epithelium

Based on the finding that PAK4-NT is responsible for interaction with Afadin, we examined whether PAK4-NT competes with endogenous PAK4 for binding Afadin and localizing to AJs. We first established a Ctrl MDCK II cell clone stably expressing GFP-PAK4-NT. In these cells, GFP-PAK4-NT accumulated at multicellular vertices as well as cell boundaries (Fig. S4I). Mixed culture of these cells with Ctrl cells showed that the endogenous full-length PAK4 was no longer localized at cell-cell boundaries in the GFP-PAK4-NT-expressing cells. This suggests that GFP-PAK4-NT can be used as a tool to interfere the localization of endogenous PAK4 at AJs (Fig. S4J), although we cannot exclude the possibility that PAK4-NT prevents PAK4 localization through different mechanisms, such as preventing interaction with active Cdc42 or other binding partners.

We then tested the effects of expressing PAK4-NT (Fig. 7A) on vertex remodeling in the *Xenopus* epithelium. Inhibiting the recruitment of endogenous PAK4 to junctions led to an increased number of higher-order vertices (Fig. 7B). We quantified the ratio of the number of tricellular vertices, 4-way vertices, or rosettes to the total number of cell vertices in control and PAK4-NT embryos. Notably, PAK4-NT expression resulted in a significant increase in ratio of 4-way vertices compared to control (Fig. 7C). Likewise, a significant increase in rosettes was observed in PAK4-NT embryos (Fig. 7C). We conclude that inhibiting endogenous PAK4 recruitment to junctions via expression of the PAK4-NT fragment prevents multicellular vertex remodeling, leading to an increased proportion of higher-order vertices in the developing *Xenopus* epithelium.

**Figure 7.**
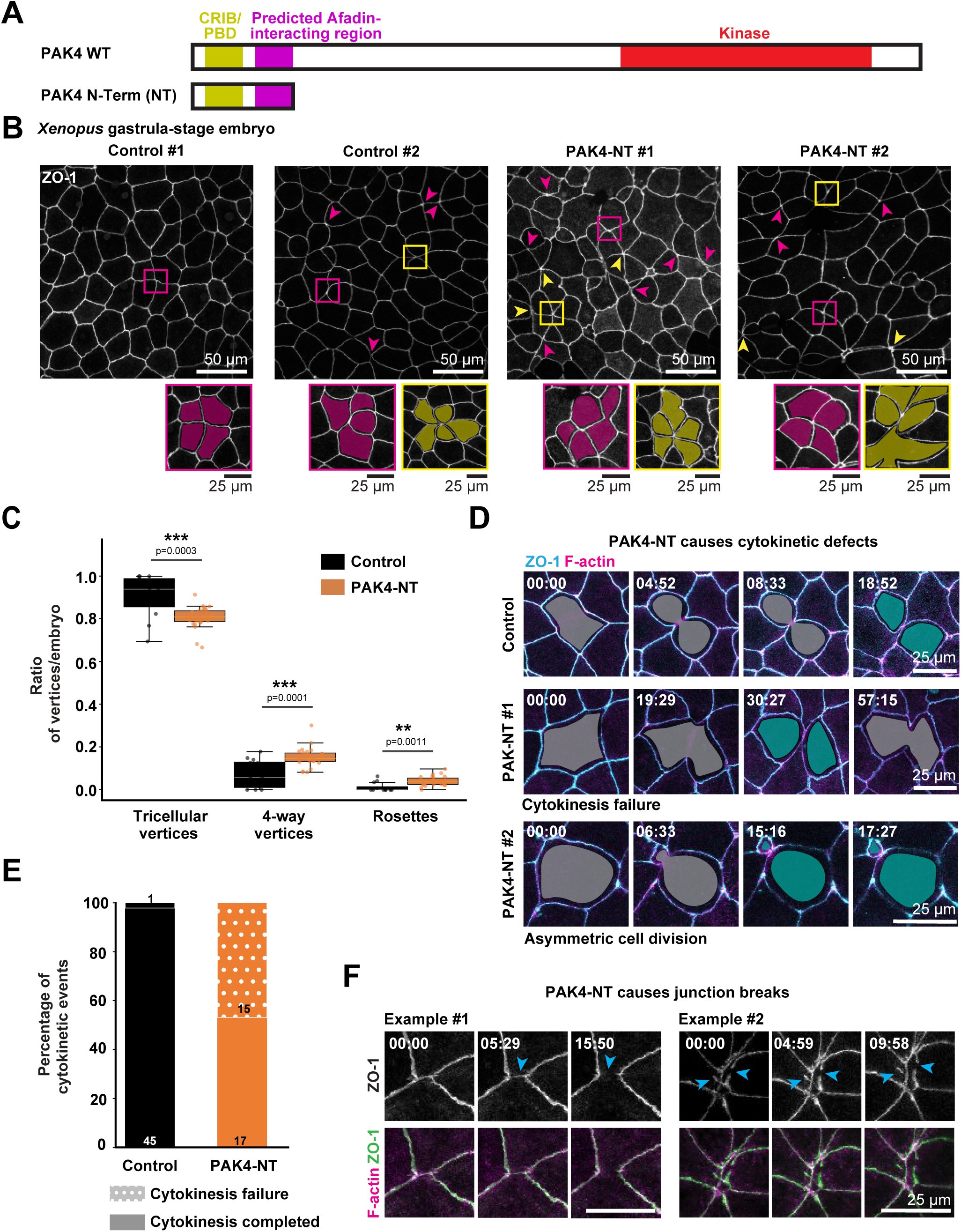
Inhibition of PAK4 kinase activity compromises junctional integrity in *Xenopus* epithelia. (A) Domain map of PAK4. N-terminal fragment of PAK4 (PAK4-NT) was expressed in the embryos to act as a dominant-negative. (B) Live confocal images of epithelial cells in the animal hemisphere of gastrula-stage *Xenopus* embryos expressing TagBFP2-ZO-1 (ZO-1) +/- PAK4-NT. Zoomed-in panels show examples highlighting 4-way (magenta) and higher-order (5 way or more) (yellow) vertices. Scale bars, 50 µm (upper), 25µm (lower). (C) Quantification of 3-way, 4-way, and higher-order vertices (control: 12 embryos; PAK4-NT: 25 embryos; 3 independent experiments). Data were compared using a two-tailed unpaired Student’s *t*-test (***, p < 0.001; **, p < 0.01) (D) Montage of control and PAK4-NT cells expressing TagBFP2-ZO-1 (ZO-1) and LifeAct-miRFP (F-actin), showing changes before and during ingression (gray) and post-cytokinesis (green). Scale bar, 25 µm. (E) Quantification of cytokinetic failure (controls: 46 cell division events from 5 embryos; PAK4-NT: 32 cell division events from 8 embryo; 3 independent experiments). (F) Montages of control and PAK4-NT embryos expressing TagBFP2-ZO-1 (ZO-1) and LifeAct-miRFP (F-actin), showing junctional discontinuities (blue arrowheads). Scale bars, 25 µm.

Live imaging of PAK4-NT-expressing embryos revealed that inhibiting PAK4 junctional localization also caused defects in epithelial cytokinesis. First, the majority of PAK4-NT cell divisions formed a 4-way vertex. In contrast to controls, where these 4-way vertices remodel into daughter-daughter or neighbor-neighbor interfaces (Fig. 4D), the neighboring cells in PAK4-NT embryos often retracted, leading to re-fusion of the daughter cells and ultimately failed cytokinesis (Fig. 7D and Video 3). Even in cases where cytokinesis was able to complete, other defects like asymmetric cell division were observed (Fig. 7D). Quantification of cytokinesis completion showed that in PAK4-NT-expressing embryos, 46.88% of cytokinesis events failed, whereas only 2.17% failed in controls (Fig. 7E). Additionally, inhibiting PAK4 recruitment to junctions via expression of PAK4-NT also caused junctional discontinuities (Fig. 7F and Video 3).

Taken together, we show that inhibition of endogenous PAK4 recruitment to junctions causes failure in multicellular vertex remodeling and cytokinesis, as well as junctional discontinuities in the *Xenopus* epithelium.

### PAK4 and Afadin are required for establishment of the epithelial barrier

Loss of epithelial integrity often results in impaired barrier function. To investigate the roles of PAK4 and Afadin in epithelial barrier maintenance, we measured transepithelial electrical resistance (TER) of *PAK4*-KO and *Afdn*-KO MDCK II cells cultured on transwell filters. Since the Ctrl cells are *Cldn2*-KO MDCK II cells, in which cation-permeable *claudin-2* was knocked out, the TER of Ctrl cells reaches 3.9-4.1 kΩ•cm^2^ on day 7 after seeding (Fig. 8A). In contrast, *PAK4*-KO cells exhibited impaired barrier function (1.5-2.0 kΩ•cm^2^) on day 7 (Fig. 8B), indicating that PAK4 is required for establishment of a tight barrier. Furthermore, the TER values of *Afdn*-KO cells remain even lower (0.06-0.21 Ω•cm^2^) than those of *PAK4*-KO cells (Fig. 8C), suggesting that Afadin contributes to the establishment of epithelial barrier not only by recruiting PAK4 to the junctions and vertices, but also through other molecular mechanism(s).

**Figure 8.**
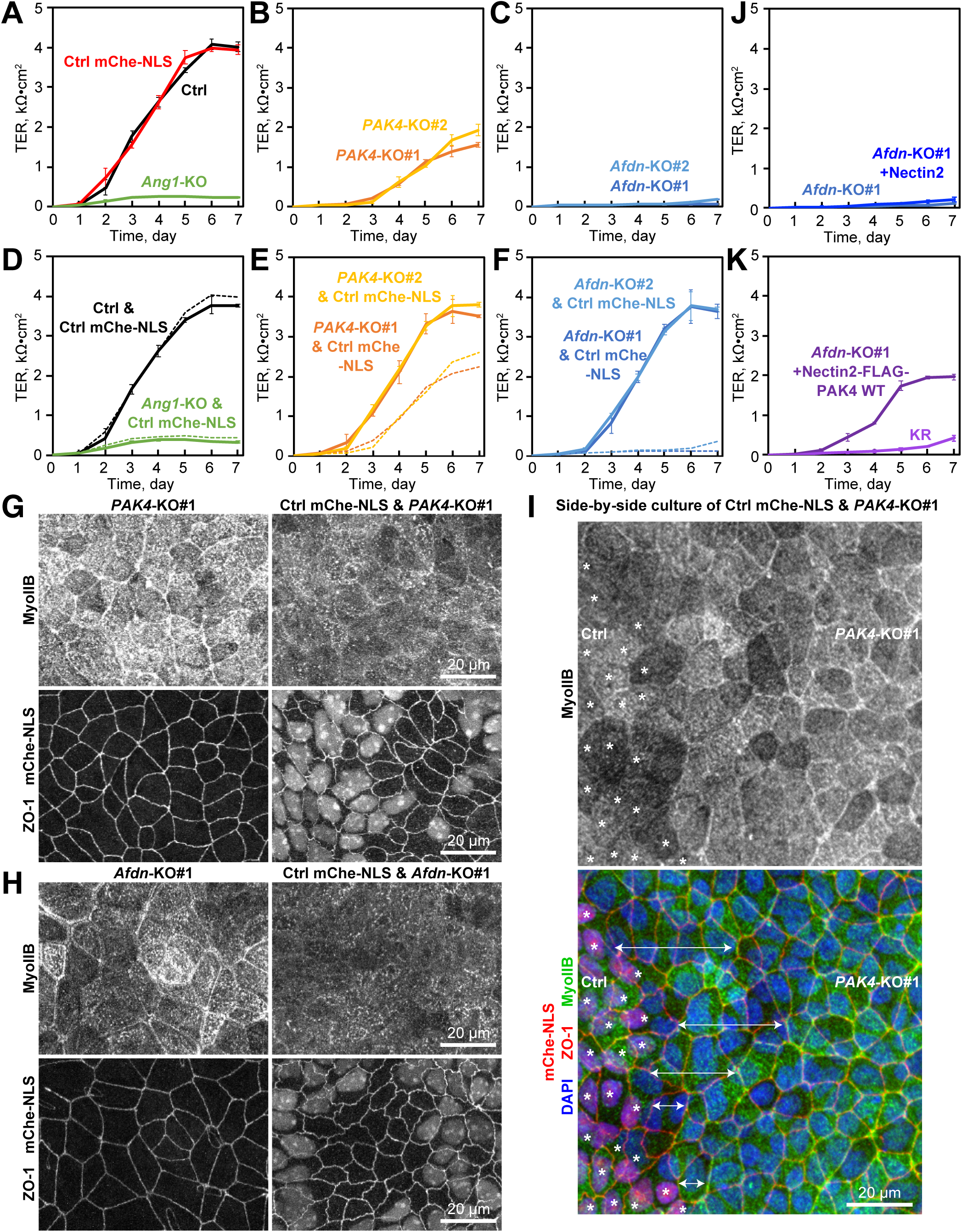
PAK4 and Afadin contribute to epithelial barrier function in cultured cells. (A-H,J-K) TER measurements for the following cell types: (A) Ctrl, Ctrl cells expressing mCherry-NLS (mChe-NLS), and *Ang1*-KO cells;(B) *PAK4*-KO cells; (C) *Afdn*-KO cells; (D) a mixed culture of mChe-NLS and *Ang1*-KO cells; (E) a mixed culture of mChe-NLS and *PAK4*-KO cells; (F) a mixed culture of mChe-NLS and *Afdn*-KO cells; (J) *Afdn*-KO #1 cells expressing Nectin2-FLAG; (K) *Afdn*-KO #1 cells expressing Nectin2-FLAG-PAK4 WT or kinase-dead (KR) mutant. Values are shown as mean ± SD. In D-F, solid lines represent TER values measured in mixed cultures, while dotted lines indicate estimated combined resistance values calculated from the separately measured TER values of mChe-NLS and KO cell cultures. (G-H) Immunofluorescence staining of the *PAK4*-KO #1 (G) or *Afdn*-KO #1 (H) cells cultured with (lower) or without (upper) mChe-NLS cells on the filter using anti-MyoIIB pAb (left) and anti-ZO-1 mAb (right). Scale bars, 20 µm. (I) Immunofluorescence staining at the boundary between mChe-NLS (asterisks) and *PAK4*-KO #1 cells grown side by side on a filter using anti-MyoIIB pAb (green), anti-ZO-1 mAb (red), and DAPI (blue). Double arrows indicate the distance between Ctrl cells and sarcomere-like MyoII structure in the *PAK4*-KO cells. Scale bar, 20 µm.

To examine whether PAK4 and Afadin function in a cell-autonomous manner, we cultured Ctrl and KO cells together and evaluated the barrier function of mixed cultures. To distinguish Ctrl and KO cells, we used a cell clone of Ctrl cells stably expressing mCherry conjugated with nuclear localization signals (NLS) (Ctrl mChe-NLS). When two cell clones with different barrier properties are mixed together and each cell clone does not affect the barrier function of the other clone, the combined TER can be predicted by harmonic mean of TER values of the two cell clones cultured separately (R_pred_ = 2 / ((1 / R_clone1_) + (1 / R_clone2_)). For example, the *angulin-1*-knockout cell clone (*Ang1*-KO) has leaky barrier properties (0.22-0.25 kΩ•cm^2^) (Fig. 8A). The TER of a mixed culture of Ctrl mChe-NLS and *Ang1*-KO cells on day 7 was 0.31-0.35 kΩ•cm^2^, which was close to the predicted value (0.44 kΩ•cm^2^) (Fig. 8D). In contrast, when *PAK4*-KO cells were mixed with Ctrl mChe-NLS cells, the TER increased much higher (3.5-3.9 kΩ•cm^2^) than the predicted values (2.2 kΩ•cm^2^ [#1] and 2.6 kΩ•cm^2^ [#2]) and reached a level almost comparable with Ctrl cells (Fig. 8E). The discrepancy was more evident in a mixed culture of Ctrl mChe-NLS and *Afdn*-KO cells (measured, 3.5-3.9 kΩ•cm^2^; predicted, 0.14 kΩ•cm^2^ [#1] and 0.37 kΩ•cm^2^ [#2]). Again, the TER of a mixed culture of Ctrl mChe-NLS and *Afdn*-KO cells was similar to that of Ctrl cells (Fig. 8F). We confirmed that the ratio of *PAK4*-KO or *Afdn*-KO cells to Ctrl mChe-NLS cells remained 1:1 throughout the experimental period (Fig. S3I), indicating that KO cells are not eliminated by mChe-NLS cells. Of interest, the sarcomere-like structure of MyoIIB in the *PAK4*-KO and *Afdn*-KO cells was not observed in the KO cells within the mixed culture (Fig. 8G-H), suggesting that neighboring Ctrl cells may convey a mechanical or chemical signal to *PAK4*-KO or *Afdn*-KO cells and suppress the formation of the sarcomere-like MyoII structure. To examine how far the suppressive effect of Ctrl mChe-NLS cells on myosin assembly extends into the neighboring *PAK4*-KO clone, we co-cultured Ctrl mChe-NLS and *PAK4*-KO cells side by side and analyzed the boundary region. The suppression of the sarcomere-like structure of MyoIIB was restricted to the first two layers of PAK4-KO cells adjacent to Ctrl mChe-NLS cells (Fig. 8I).

Since artificial targeting of PAK4 to junctions using the Nectin2-FLAG-PAK4 construct partially rescued the epithelial integrity defects, we speculated that artificial PAK4 targeting could also improve the barrier properties of *Afdn*-KO cells. Expressing Nectin2 alone in *Afdn*-KO cells did not affect barrier function (Fig. 8J). As predicted, *Afdn*-KO cells stably expressing Nectin2-FLAG-PAK4 exhibited much higher TER values compared to the parental *Afdn*-KO cell clone (Fig. 8K), although the barrier function was not fully rescued to the Ctrl level. The kinase-dead version (Nectin2-FLAG-PAK4 K351R) failed to rescue the barrier defect (Fig. 8K), indicating that kinase activity of PAK4 is important to support barrier function.

These data suggest that PAK4 recruitment by Afadin is important for the establishment of the epithelial barrier. They also suggest that Ctrl NLS-mChe cells affect the cell structure and mechanical properties of neighboring *PAK4*-KO and *Afdn*-KO cells and can rescue the barrier defects of KO cells.

### PAK4 inhibition disrupts epithelial barrier function in *Xenopus* embryos

In PAK4 inhibitor-treated embryos, small openings were regularly observed at multicellular rosettes (Fig. S5A). Thus, we wanted to investigate the barrier function of the *Xenopus* epithelium upon PAK4 inhibition. To assess changes in barrier function, we used the Zinc-Based Ultrasensitive Microscopic Barrier Assay (ZnUMBA) to visualize dynamic barrier leaks, which appear as increases in the fluorescence intensity of a zinc ion indicator, FluoZin3 (Stephenson et al., 2019; Higashi et al., 2023b) (Fig. 9A). Whole-field and junctional FluoZin3 intensity were not significantly different in control vs. PAK4 inhibitor-treated embryos (Fig. S5B-C), indicating that overall barrier function is not severely affected. In controls, there were occasional local barrier leaks (increases FluoZin3 signal) (Fig. 9B). In contrast, PAK4 inhibitor-treated embryos exhibited increased leaks – particularly at vertices (Fig. 9B and Video 4). Indeed, PAK4 inhibitor-treated embryos had a significantly higher percentage of barrier leaks at vertices (32.74±4.51%) compared to controls (16.83±1.84%) (Fig. 9C). Additionally, PAK4 inhibitor-treated embryos showed a significant increase in FluoZin3 intensity at higher-order vertices (4-way or higher) compared to controls (Fig. 9D). Of note, the barrier leaks observed in controls were quickly repaired, and FluoZin3 signal returned to baseline (Fig. 9E). In contrast, in PAK4 inhibitor-treated embryos, repeated or persistent leaks were observed at some vertices (Fig. 9E).

**Figure 9.**
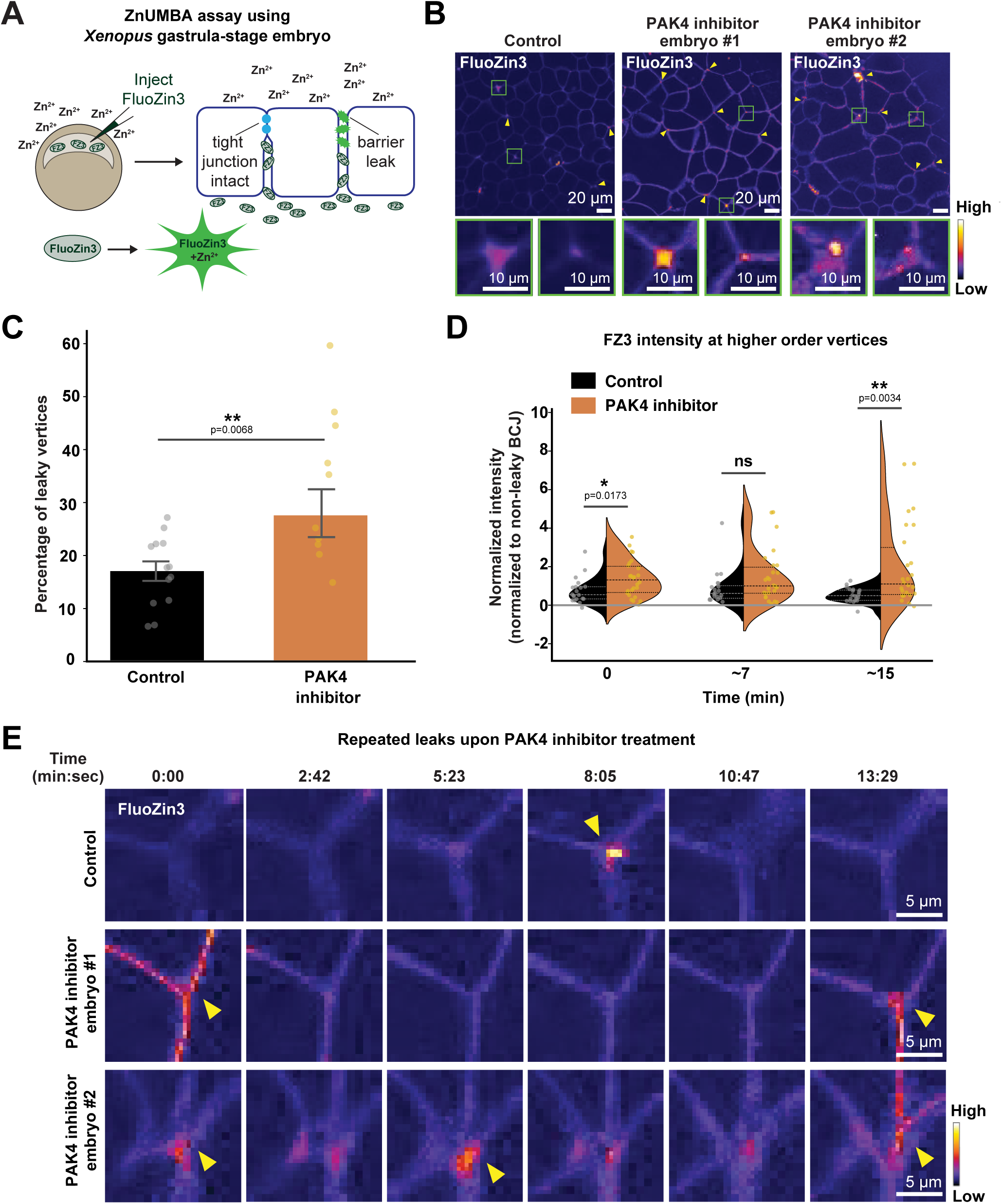
PAK4 inhibition impairs epithelial barrier integrity at higher-order vertices in *Xenopus* embryos. (A) Schematic of the ZnUMBA assay. Disruption of TJ allows Zn^2+^ to bind FluoZin-3, resulting in increased FluoZin-3 fluorescence, which indicates barrier leakage. (B) Live confocal images of FluoZin-3 signal (FIRE LUT) in control and PAK4 inhibiter-treated *Xenopus* embryos. Yellow arrowheads indicate increased signal at higher-order vertices. Zoomed-in panels (green boxes) highlight barrier leaks at higher-order vertices. Scale bars, 20 µm (upper), 10 µm (lower). (C) Quantification of leaky vertices (control: 13 embryos, 1375 vertices; PAK4 inhibitor: 10 embryos, 796 vertices; 3 independent experiments). Error bars, SEM. Data were compared using a two-tailed unpaired Welch’s *t*-test (**, p < 0.01). (D) Quantification of FluoZin-3 intensity at higher-order vertices (4-way and above), normalized to non-leaky bicellular junctions at 0, 7, and 15 min after the onset of imaging (control: 12 embryos, 20 vertices; PAK4 inhibitor: 11 embryos, 26 vertices; 3 independent experiments). Violin plots show the median (solid line) and 25th/75th percentiles (dotted lines), with individual measurements overlaid (circles). Data are compared using a two-tailed unpaired Student’s *t*-test (*, p < 0.05, **, p < 0.01). (E) Montage of FluoZin-3 signal (FIRE LUT) at representative higher-order vertices in control and PAK4 inhibitor-treated embryos, showing leaks (yellow arrowheads). Scale bars, 5 µm.

Next, we wanted to test whether inhibiting endogenous PAK4 recruitment to junctions via expressing PAK4-NT would disrupt epithelial barrier function. In control embryos, baseline levels of FluoZin3 signal at cell-cell junctions were low, with occasional local barrier leaks that were rapidly repaired, indicating that barrier function is intact (Fig. 10A). In contrast, PAK4-NT-expressing embryos exhibited striking barrier defects. PAK4-NT embryos had a higher number of leaks specifically localized to the multicellular vertices (Fig. 10A). In 50% of PAK4-NT-expressing embryos, the whole-field intensity of FluoZin-3 increased over time (Fig. 10A-B), indicating global barrier disruption (Video 5). Significant differences between control and PAK4-NT-expressing embryos were observed for both whole-field and junctional FluoZin3 intensity (Fig. 10C-D). Additionally, PAK4-NT-expressing embryos exhibited a significantly higher percentage of leaks at vertices (63.85±9.23%) compared to controls (16.83±1.84%) (Fig. 10E). Quantification of the intensity of FluoZin3 signal at higher-order vertices (4-way or higher) showed that PAK4-NT-expressing embryos exhibited a significant increase in FluoZin3 intensity at higher-order vertices compared to controls (Fig. 10F). The barrier leaks observed in controls were rapidly repaired (Fig. 10G). However, PAK4 N-term-expressing embryos, often failed to effectively repair their leaks, and persistent or repeated leaks were common (Fig. 10G). The global barrier disruption observed in these embryos could be a result of the failure to repair local leaks.

**Figure 10.**
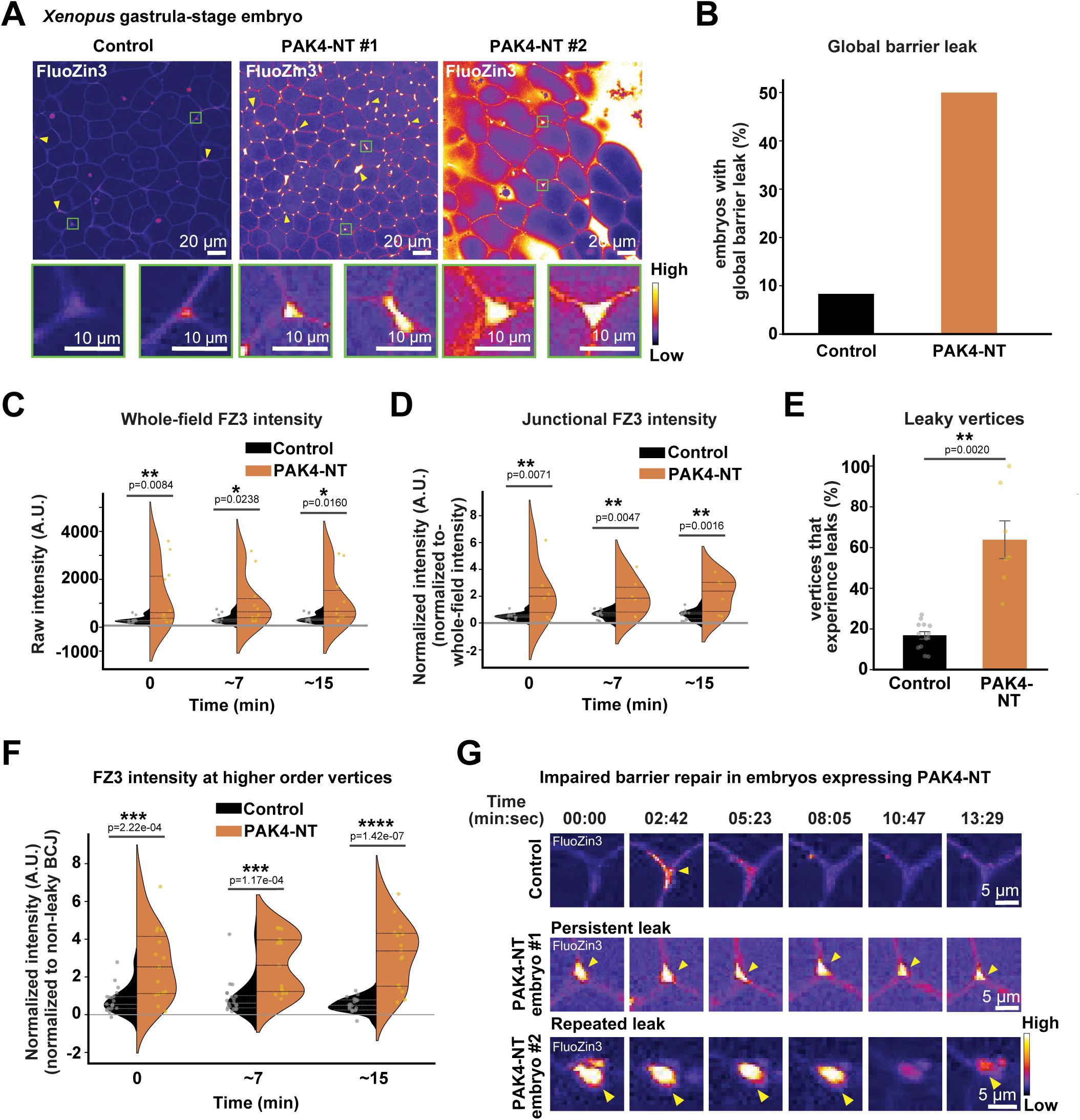
PAK4-NT impairs epithelial barrier function at higher-order vertices in *Xenopus* embryos. (A) Live confocal images of FluoZin-3 signal (FIRE LUT) in control and PAK4-NT-expressing *Xenopus* embryos. Yellow arrowheads indicate elevated signal at higher-order vertices. Zoomed-in panels (green boxes) highlight local barrier leaks. Scale bars, 20 µm (upper), 10 µm (lower). (B) Percentage of embryos exhibiting global barrier leaks. n = 12 (control, black), 10 (PAK4-NT, orange). (C) Quantification of whole-field FluoZin-3 intensity at 0, ∼7, and ∼15 min after the onset of imaging (control: 13 embryos; PAK4-NT: 10 embryos; 3 independent experiments). (D) Quantification of junctional FluoZin-3 intensity at 0, ∼7, and ∼15 min after the onset of imaging (control: 13 embryos, PAK4-NT: 6 embryos; 3 independent experiments). (E) Quantification of vertices exhibiting barrier leaks (control: 13 embryos, 1375 vertices; PAK4-NT: 7 embryos, 815 vertices; 3 independent experiments). (F) Quantification of FluoZin-3 intensity at higher-order vertices (4-way and above), normalized to the non-leaky bicellular junctions (control: 12 embryos, 20 vertices; PAK4-NT: 6 embryos, 15 vertices; 3 independent experiments). (G) Montage of FluoZin-3 signal (FIRE-LUT) at representative higher-order vertices showing repeated and persistent leaks (yellow arrowheads) in PAK4-NT-expressing embryos. Scale bars, 5 µm. For violin plots (C, D, F), solid and dotted lines indicate the median and the 25th/75th percentiles. Data were analyzed using a two-tailed unpaired Student’s *t*-test (C, D, F) or Welch’s *t*-test (E). *, p < 0.05; **, p < 0.01; ***, p < 0.001; ****, p < 0.0001.

Taken together, these results suggest that PAK4 is required for maintaining epithelial barrier function *in vivo*. Pharmacological inhibition of PAK4 or expression of PAK4-NT causes increased leaks – both in number and severity – at multicellular vertices, challenging the barrier integrity of the developing epithelium.

## Discussion

### Vertex remodeling plays a fundamental role in epithelial tissue organization and dynamics

Vertex remodeling drives directional cell rearrangements, contributing to processes like convergent extension during development in multicellular organisms. In adult tissues like the intestine, similar vertex remodeling events are believed to occur during the steady-state renewal of epithelial sheets, where cells derived from stem cell niches migrate across the epithelial plane. In both contexts, vertex remodeling must meet two seemingly opposing requirements: 1) maintaining epithelial integrity and barrier function at tTJs and 2) enabling flexible reorganization of cell neighbors. Our study demonstrates that PAK4 helps fulfill both of these roles during passive vertex remodeling events, as shown by gene knockouts in cultured MCDK II epithelial monolayers and live imaging of the *Xenopus* embryonic epithelium. However, whether PAK4 is also required for active, developmentally-driven remodeling during convergent extension remains an open question. Of interest, improper folding of the caudal portion of the neural tube in PAK4-null mouse embryos (Qu et al., 2003) suggests a role for PAK4 in convergent extension.

### PAK4 localizes to remodeling vertices and is involved in the resolution of higher-order vertices

Vertex remodeling at higher-order vertices can be divided into three stages: contraction of bicellular junctions leading to vertex convergence, formation of multicellular vertices in which four or more cells share a vertex, and resolution of such multicellular vertices into new configuration of lower-order vertices. We observed that PAK4 preferentially localizes to vertices undergoing remodeling, particularly to 4-way vertices and rosettes, and that PAK4 knockout or inhibition leads to a significant accumulation of unresolved rosettes. These findings suggest that PAK4 is primarily involved in the resolution phase. While Afadin is required for the recruitment of PAK4 to junctions, the vertex- and rosette-specific enrichment of PAK4 is more prominent than that of Afadin, implying that additional signals are needed. Candidates include local activation of Cdc42 (Oda et al., 2014), phosphorylation of Afadin (Yu and Zallen, 2020), and liquid-liquid phase separation (LLPS)-mediated condensate formation by Afadin (Kuno et al., 2025). Previous studies have identified additional regulators such as Sidekick (Lye et al., 2014; Finegan et al., 2019; Letizia et al., 2019; Uechi and Kuranaga, 2019; Malin et al., 2022), M6 (Wittek et al., 2020; Esmangart de Bournonville and Le Borgne, 2020; Ikawa et al., 2023), PTEN (Bardet et al., 2013; Malin et al., 2024), Ajuba (Razzell et al., 2018; Ikawa et al., 2023), and Vinculin (van den Goor et al., 2024; Landino et al., 2025) as modulators of vertex dynamics. Interestingly, PAK4 was reported to interact with Vinculin in the context of podosome formation in myeloid cells (Foxall et al., 2019), suggesting a possible link between mechanical remodeling and actin-cytoskeleton regulation. Recently, it was reported that the *Drosophila* homolog of PAK4, mbt, regulates Sidekick localization at tAJs, raising a possibility that PAK4 regulates the structure of tAJs through Sidekick (Gandhi et al., 2025), although it is still unclear whether the vertebrate homologs of Sidekick, Sdk1 and Sdk2, are involved in tAJ architecture and function. Whether such interactions also cooperate at vertebrate epithelial vertices, and how they contribute to rosette resolution and junctional rearrangement, are important topics for future investigation.

### PAK4’s kinase activity is essential for its function in vertex remodeling

Vertex remodeling was impaired by pharmacological inhibition of PAK4 (Figs. 3 and 4) as well as by genetic ablation of PAK4 (Fig. 5), and this phenotype was not rescued by expression of a kinase-dead mutant (Fig. S2), highlighting the importance of its kinase activity in this process. However, the direct phosphorylation targets of PAK4 in this context remain unknown. It was reported that U2OS cells treated with a PAK4 inhibitor did not exhibit changes in Afadin phosphorylation, but several candidate proteins proximal to PAK4 were identified, including cingulin, p120-catenin, DLG5, Scrib, ARHGEF11 (also known as PDZ-Rho GEF), and ZO-1/2 (Baskaran et al., 2021). Some of these proteins may act downstream of PAK4 to regulate membrane plasticity, cell polarity, and cytoskeletal reorganization during rosette resolution. Notably, ARHGEF11 binds to PAK4 (Barac et al., 2004) and has been reported as a phosphorylation target of PAK4 in invadosome regulation in cancer cells (Nicholas et al., 2016), raising the possibility of shared pathways in epithelial and invasive structures. PAK4 is known to undergo autophosphorylation at K350 (K351 in dog PAK4) within the activation loop of the kinase domain, and this modification is required for its activation (Abo et al., 1998). Although this site is generally considered constitutively phosphorylated (Callow et al., 2002), long-term kinase inhibition or a loss-of-function mutation may disrupt this phosphorylation and thereby alter the conformation of PAK4. In this scenario, PAK4 may act not only as a kinase but also as a structural scaffold at remodeling vertices, influencing junctional and cytoskeletal architecture. Supporting this view, we observed repositioning of PAK4-localizing vertices during cell rearrangements (Fig. 2F-G), suggesting a transient shift in junctional material properties from solid-like to liquid-like states. LLPS-prone proteins, such as Afadin (Kuno et al., 2025), are promising candidates for mediating such transitions.

### Epithelial integrity maintenance by Afadin and recruitment of PAK4

Afadin has been recognized as a key component in stabilizing actin filaments at AJs through its interactions with nectins and α-catenin (Mandai et al., 1997; Takahashi et al., 1999; Tachibana et al., 2000; Pokutta et al., 2002; Sakakibara et al., 2020), a mechanism that is central to the maintenance of epithelial integrity. Building on this framework, our study uncovers an additional function: Afadin recruits PAK4 to cell-cell junctions where it is important for maintaining epithelial integrity. This is evidenced by the findings that PAK4 and Afadin interact (Fig. S4), and artificial targeting of PAK4 to cell-cell junctions in *Afdn*-KO cells partially rescued epithelial integrity defects (Fig. 6E-I). The incomplete rescue strongly suggests that Afadin contributes to epithelial stability through mechanisms beyond PAK4 recruitment, in line with its canonical role in actin anchoring (Mandai et al., 1997; Sawyer et al., 2009; Sakakibara et al., 2020; Gong et al., 2025; Jensen et al., 2025). A recent report further refines this view by indicating that mature AJs (named *zonula adherens matura*) can be partitioned into two belt-like zones: and apical Afadin/nectin-populated zone and a basal cadherin/catenin-populated zone, with F-actin preferentially tethered at the Afadin/nectin zone (Mangeol et al., 2024). The configuration thought to be important for assembling AJs capable of withstanding mechanical tension. At the same time, Afadin is also reported to cooperate with α-catenin and Vinculin to stabilize actin bundles (Gong et al., 2025), implying crosstalk between Afadin/nectin- and cadherin/catenin-based zones. How Afadin-recruited PAK4 interfaces with this zonal architecture and with classical actin regulators remain unclear, although our data suggest that PAK4 itself localizes to the Afadin/nectin-based zone (Fig. 2E). Resolving these relationships will be an important direction for future work.

The N-termini of both Afadin and its *Drosophila* homolog Cno harbor tandem RA domains (RA1 and RA2), which have been regarded as modules for binding Ras family GTPases (Kuriyama et al., 1996; Boettner et al., 2000) such as Rap1. This model has explained the genetic interaction and overlapping functions of Afadin/Cno and Rap1 (Boettner et al., 2003; Sawyer et al., 2009; Choi et al., 2013). However, a recent study revealed that the sequence of RA1, but not that of RA2, is essential for epithelial integrity (McParland et al., 2024). This difference is puzzling given that both domains are capable of binding Rap1, albeit with different affinities (Linnemann et al., 1999; Wohlgemuth et al., 2005; McParland et al., 2024). Our structural predictions suggesting that RA1 also provides a platform for recruitment of PAK4 (or its *Drosophila* homolog mbt) would explain the unique requirement of the RA1 domain.

### The role of epithelial vertices in maintaining barrier function

This study demonstrated that loss of PAK4 or inhibition of its activity leads to impaired epithelial barrier function, with the ZnUMBA assay revealing particularly pronounced barrier defects at tricellular and multicellular vertices. These sites are known to be mechanically vulnerable because they concentrate tensile forces, and the observed defects highlight the importance of PAK4 in maintaining junctional integrity under mechanical stress. At tricellular junctions within vertebrate epithelia, barrier properties are normally ensured by specialized tTJs, composed of proteins such as tricellulin and angulins (Higashi and Miller, 2017; Higashi and Chiba, 2020). Upon PAK4 inhibition, however, angulin-1 localization at multicellular vertices exhibited gap-like openings at the center (Fig. S5A), suggesting that PAK4 is essential for preserving the architecture of tTJs. Previous studies have shown that epithelial barriers are dynamically maintained by mechanisms including localized and transient Rho activation (“Rho flares”) (Stephenson et al., 2019; Varadarajan et al., 2022; Chumki et al., 2022; Craig et al., 2025) and claudin-7 polymerization following its release from EpCAM or Trop2 by proteases (Higashi et al., 2023a; Cho et al., 2025). Rho flares frequently occur at multicellular vertices (Stephenson et al., 2019), and polymerized claudin-7 also accumulates at vertices (Higashi et al., 2023a). Together, these observations indicate that multiple vertex-centered mechanisms act in concert to sustain barrier function and that PAK4 likely interfaces with these repair systems. Elucidating the phosphorylation targets of PAK4 will therefore provide important insight into the molecular pathways governing barrier maintenance at tricellular vertices.

### New insights into the sarcomere-like structure at bicellular cell-cell junctions

In *PAK4*-KO and *Afdn*-KO cells, we observed that myosin IIA and IIB exhibit a punctate, sarcomere-like organization along cell-cell junctions. Although the molecular mechanism underlying the generation of these sarcomere-like arrays remains unclear, their formation does not require either PAK4 or Afadin. Mixed-culture experiments with Ctrl cells demonstrated that these proteins do not suppress sarcomere-like structure formation through a direct, cell-autonomous mechanism but rather influence junctional actomyosin patterning in a cell-nonautonomous manner. This phenotype may reflect an imbalance in mechanical forces acting at cell-cell junctions, where tensile forces perpendicular and parallel to the junction axis normally cooperate to support epithelial morphology and mechanoresponsiveness (Lecuit and Yap, 2015; Tang, 2018; Angulo-Urarte et al., 2020; Lenne et al., 2021). Among these, perpendicular forces, in particular, drive the zigzag morphology of cell-cell junctions (Miyazaki et al., 2023), which is prominent in Ctrl MDCK II cells. Attenuation of perpendicular forces may allow parallel actomyosin arrays to stabilize, thereby promoting sarcomere-like patterning along bicellular junctions and increasing tension at vertices. This raises the question of which molecules are responsible for generating perpendicular forces at cell-cell junctions. ZO proteins are prime candidates, as ZO-1/ZO-2 double depletion induces sarcomere-like structures (Yamazaki et al., 2008; Fanning et al., 2012; Choi et al., 2016; Otani et al., 2019), whereas ZO-1 overexpression exaggerates junctional curvature (Tokuda et al., 2014). ZO-1 may exert these perpendicular forces through mechanosensitive conformational changes (Spadaro et al., 2017), which are reduced in *claudins/JAM-A*-KO cells that also display junctional myosin bundles (Nguyen et al., 2024). Notably, both ZO-deficient cells and Afadin-deficient cells failed to develop sarcomere-like myosin arrays when surrounded by control cells (Choi et al., 2016) (Fig. 8H), implicating a similar cell-nonautonomous mechanism. For example, Afadin and PAK4 might regulate ZO-1 conformation similar to claudins/JAM-A. Alternatively, they might generate perpendicular tensile forces on bicellular junctions independently of ZO proteins, although this possibility appears less likely, since junctions are straightened in ZO-1/2 double-knockdown cells despite the increased junctional accumulation of Afadin (Choi et al., 2016), suggesting that perpendicular force generation does not simply correlate with the amount of Afadin at junctions. Of interest, the *Drosophila* homolog of ZO-1, Polychaetoid, also works in parallel with Afadin homolog, Canoe, to maintain epithelial integrity during development (Manning et al., 2019). Future studies should address how Afadin/PAK4 and ZO proteins cooperatively maintain epithelial integrity.

### Concluding remarks

Here, we demonstrate a novel role for PAK4 and Afadin in the maintenance of epithelial integrity and barrier function during vertex remodeling. Defining phosphorylation targets of PAK4 and downstream signaling pathways is an important future direction.

## Materials and methods

### Cell culture

MDCK II and HEK293T cells were cultured in Dulbecco’s Modified Eagle Medium (DMEM; D5796; Sigma-Aldrich, St. Louis, MO, USA) supplemented with 5% fetal bovine serum (FBS; F7524; Sigma-Aldrich). Cells were maintained in a 5% CO_2_ incubator at 37°C. All KO clones of MDCK II cells were established from a parental *cldn2*-KO MDCK II cell clone (Saito et al., 2021). Thus, the *cldn2*-KO cell line is referred to as the “Ctrl” cell line throughout this manuscript. The *cldn2*-KO cell clone expressing mCherry conjugated with 3xnuclear localization signal (mChe-NLS) was described previously (Higashi et al., 2023b).

### Antibodies used in cell culture

The following primary antibodies were used: Rabbit anti-PAK4 monoclonal antibody (mAb) (clone E7H7R; #52694), rabbit anti-phospho-Myosin light chain 2 (Thr18/Ser19) polyclonal antibody (pAb) (#3674), and rabbit anti-EpCAM mAb (clone E6V8Y; #93790) (Cell Signaling Technology, Danvers, MA, USA); rabbit anti-l/s-Afadin pAb (#A0224) and mouse anti-vinculin mAb (clone VIN-11-5; #V4505) (Sigma-Aldrich); rabbit anti-non-muscle Myosin heavy chain II-A (MyoIIA) pAb (#909801) and rabbit anti-non-muscle Myosin heavy chain II-B (MyoIIB) pAb (#909901) (BioLegend, San Diego, CA, USA); rat anti-ZO-1 (alpha+) mAb (clone R40.76; sc-33725), mouse anti-claudin-1 mAb (clone XX7; sc-81796), and mouse anti-beta-catenin mAb (clone E-5; sc-7963) (Santa Cruz Biotechnology, Dallas, TX, USA); mouse anti-DYKDDDDK tag (FLAG-tag) mAb (clone 1E6; #012-22348) (Wako-Fujifilm, Osaka, Japan); mouse anti-p120-catenin (CTNND1) mAb (clone 2F7H8; #66208-1-Ig), mouse anti-GFP tag mAb (clone 1E10H7; #66002-1-Ig), and rabbit anti-mCherry pAb (#26765-1-AP) (Proteintech, Rosemont, IL, USA); rabbit anti-claudin-3 pAb (#34-1700), mouse anti-claudin-4 mAb (clone 3E2C1; #32-9400), and Alexa Flour 488-conjugated mouse anti-ZO-1 mAb (Clone ZO1-1A12; #339188) (Thermo Fisher Scientific, Waltham, MA, USA); rabbit anti-claudin-7 pAb (#18875) (Immuno-Biological Laboratories, Fujioka, Gunma, Japan); rabbit anti-Partitioning-defective 3 (Par3) pAb (#07-330) (Merck Millipore, Burlington, MA, USA).

For immunofluorescence imaging, the following secondary antibodies were used: Alexa Fluor 488-conjugated AffiniPure donkey anti-mouse IgG (H+L) pAb (#715-545-151), Alexa Fluor 488-conjugated AffiniPure donkey anti-rabbit IgG (H+L) pAb (#711-545-152), Alexa Fluor 488-conjugated AffiniPure donkey anti-rat IgG (H+L) pAb (#712-545-153), Cy3-conjugated AffiniPure donkey anti-mouse IgG (H+L) pAb (#715-165-151), Cy3-conjugated AffiniPure donkey anti-rabbit IgG (H+L) pAb (#711-165-152), Cy3-conjugated AffiniPure donkey anti-rat IgG (H+L) pAb (#712-165-153), Alexa Fluor 647-conjugated AffiniPure donkey anti-rabbit IgG (H+L) pAb (#711-605-152), and Alexa Fluor 647-conjugated AffiniPure donkey anti-mouse IgG (H+L) pAb (#715-605-151) (Jackson ImmunoResearch Laboratories, West Grove, PA, USA).

For immunoblotting, the following secondary antibodies were used: Horseradish peroxidase (HRP)-linked sheep anti-mouse IgG pAb (#NA931V) (GE Healthcare, Chicago, IL, USA); HRP-linked goat anti-rabbit IgG pAb (#7074P) (Cell Signaling Technology); HRP-linked anti-β-actin mAb (clone C4; sc-47778HRP; Santa Cruz Biotechnology).

### Generation of KO MDCK II cell lines

*Cldn2*-KO cells were described previously (Saito et al., 2021). To generate *PAK4*-KO and *Afdn*-KO cells, DNA oligonucleotides encoding single-guide RNAs (sgRNAs) were synthesized (Integrated DNA Technologies; Coralville, IA, USA), and the annealed DNA oligonucleotides were cloned into pSpCas9 (BB)-2A-Puro (PX459) plasmids (plasmid #62988; Addgene, MA, USA) (Ran et al., 2013) at the *Bbs I* cleavage sites. Two gRNAs were designed for each gene to induce an excision of exons encoding the entire coding region (*PAK4*) or the initiation codon (*Afdn*) (Saito et al., 2024). Parental cells were transiently transfected with a pair of plasmids encoding sgRNAs using PEI-max (Polysciences, PA, USA) and selected using 3 µg/mL of puromycin (Sigma-Aldrich) for 1 d. The cells were then sparsely seeded onto 10-cm dishes and cell clones were obtained by scraping off the colonies. Genomic DNA was extracted from cell clones and screened by PCR using GoTaq DNA polymerase (#M7123; Promega, Madison, WI, USA) and specific primers. PCR products of KO clones were sequenced (Macrogen, Tokyo, Japan) to determine the precise deletion sites. After the establishment of cell clones, we confirmed that the cells were no longer resistant to puromycin.

### DNA constructs used in cell culture

A cDNA library of MDCK II cells was synthesized using the PrimeScript II 1st strand cDNA synthesis kit (#6210A; Takara Bio). cDNAs encoding canine *PAK4*, *AFADIN* (*Afdn*), and *NECTIN-2* and their fragments were amplified using PrimeSTAR GXL DNA polymerase (#R050A; Takara Bio) from the cDNA library of MDCK II cells. Primer sets used are 5’-

AT*GGATCC*GGCACCATGTTTGGGAAGAAGAAG-3’ and 5’-

AT*GAATTC*ATCTGGTGCGGTTCTGGCGC-3’ (*PAK4*); 5’-

ATG*AGATCT*GGGAGGATGTCGGCGGGCGGC-3’ and 5’-

ATT*GAATTC*ACTTGGTGTTCAGTTCGTTTTC-3’ (*AFADIN*); 5’-

AATAAT*GCTAGC*<u>ATG</u>GCCCGGGCCGCAGCCCTCC-3’ and 5’-AATAAT*GCGGCCGC*CCACATAAATCGCCCGCCGTGAAATGAGG-3’ (*NECTIN-2*). Restriction sites (BamH I and EcoR I for *PAK4*; Bgl II and EcoR I for *AFADIN*; Nhe I and Not I for *NECTIN-2*) are italicized, and the start and termination codons are underlined. Five restriction sites in the *AFADIN* cDNA sequence were disrupted by introducing a silent mutation using specific primer sets (5’-

GATGACATTGA*GAACTC*CCGACTGGCTGC-3’ and 5’-

GCAGCCAGTCGG*GAGTTC*TCAATGTCATC-3’; GGAGGATGCAG*GAGTTC*CGCAGCTCGGATG-3’ and 5’-

CCGAGCTGCG*GAACTC*CTGCATCCTCCTCTC-3’; 5’-

CGGATCAG*GAATCC*AGCCCCACCACTGC-3’ and 5’-

GTGGTGGGGCT*GGATTC*CTGATCCGAAC-3’; 5’-

GCAAGCATT*GAGTTC*AGGGAAAGTTCTG-3’ and 5’-

CTTTCCCT*GAACTC*AATGCTTGCAGGC-3’; 5’-

GCAGGACCG*AGACCT*TAGTCGAATCAC-3’ and 5’-

GTGATTCGACTA*AG<u>G</u>TCT*CGGTCCTGC-3’). The fragments were digested with restriction enzymes, purified with Nucleospin Gel and PCR Clean-up kit (#U0609C; Takara Bio), and cloned into the pCAG vector with 1xFlag (DYKDDDDK) and 2xStrep II (WSHPQFEK) tags (pCAG-FS2 vector) or pCAG-nGFP (Saito et al., 2021). The DNA sequences were verified (Macrogen, Eurofin Genomics [Tokyo, Japan], or Azenta [Burlington, MA, USA]).

### Immunofluorescence microscopy of MDCK II cells

For fluorescence microscopy of MDCK II cells, 2 x 10^5^ cells/mL of cells were seeded onto glass coverslips placed in wells of 12-well plates. In Fig. 3 experiment, PF-3758309 (#6005; Tocris) dissolved in DMSO (#D2550; Sigma-Aldrich) was added 4 hrs before fixation. After 36-44 hrs, the cells were fixed with 100% ethanol (#057-00456; Wako) for 15 min at –20°C or 2.7% formalin (#064-00406; Wako) (1% formaldehyde) in PBS for 20 min at 20-28°C and permeabilized with 0.2% Triton X-100 (#168-11805; Wako) in PBS.

For staining of established epithelial sheets of MDCK II cells, 1.0 x 10^5^ cells/mL of cells were seeded onto polycarbonate Transwell filters with 0.4-µm pore size (#3401, Corning). Culture medium was replaced daily from the day after seeding. After 6 days, the cells were fixed with 100% ethanol for 15 min at –20°C or 2.7% formalin in PBS for 20 min at 20-28°C and permeabilized with 0.2% Triton X-100 in PBS. For side-by-side culture (Fig. 8I), 80 µL suspensions of Ctrl and *PAK4*-KO#1 cells at 5.0 x 10^5^ cells/mL density were placed on each half of the Transwell filter, carefully avoiding contact between the two droplets. On the day of seeding, the plate containing filters were gentry returned to the incubator to prevent the two cell populations from merging. From the following day, once cells had adhered to the filter, the culture medium was replaced daily in the same manner as for standard culture.

Coverslips or filter membranes excised from the filter cups were washed with PBS thrice. After blocking with 2% bovine serum albumin (BSA) (#A7030; Sigma-Aldrich) in PBS, the samples were incubated with primary antibodies diluted in PBS containing 0.2% BSA for 1 h at 20-28°C. After washing with PBS, the samples were incubated with fluorescently labelled secondary antibodies diluted in PBS containing 0.2% BSA for 1 h at 20-28°C. After washing, the cells were embedded in FLUORO-GEL II with DAPI (Electron Microscopy Sciences, PA, USA). The samples were observed using a laser scanning confocal microscope (FV1000; Olympus, Tokyo, Japan) with a 60x oil-immersion objective lens (UPlanSApo 60x, Olympus) at laser wavelengths of 405, 488, 559, and 635 nm, or a fluorescence microscope (BX61, Olympus) with a 40x objective lens (UPlanSApo 40x, Olympus) equipped with a mercury lamp, dichroic filter sets (NIBA, WIG and WU) and a cooled CCD camera (DP71, Olympus). Images were acquired using the FluoView ver. 4.2b (Olympus) or CellSens ver. 1.14 (Olympus) and processed using ImageJ (NIH) and Photoshop (Adobe, CA, USA).

### Immunofluorescence microscopy of mouse small intestine

All experiments were carried out strictly adhered to the Japanese Guidelines for Proper Conduct of Animal Experiments. The protocol for the experiments using animals (#2024025) was reviewed by the Animal Care and Use Committee of Fukushima Medical University and approved by the university’s president. The 8-week-old female C57/BL6 mice were euthanized by cervical dislocation, and dissected tissue blocks were fixed in 4% paraformaldehyde (PFA) at 4°C for 2 hr, serially soaked in 15% and 30% sucrose in PBS, and embedded in OCT compound (Sakura Finetek Japan, Tokyo, Japan). The frozen tissues were cut into 6-µm-thick sections in a cryostat (Leica CM1950; Leica Biosystems, Nussloch, Germany) at –20°C, soaked in 0.5% Antigen activator Immunosaver (#333; Nisshin EM, Tokyo, Japan) solution at 70°C for overnight. The sections were permeabilized with PBS containing 0.1%(w/v) Triton X-100 for 15 min at 20-28°C, blocked with 2% BSA in PBS, stained with primary and secondary antibodies, and observed with FV1000 confocal microscope as described above.

### Quantification of fluorescence intensity at bicellular junctions and multicellular vertices in mouse tissues and MDCK II cells

Z-stacks of confocal fluorescence images were used for quantification. Fluorescence intensities were measured using ImageJ software. Circular regions of interest (ROIs) with a diameter of 5 pixels were placed manually at each vertex, which were classified as tricellular vertices or tetracellular vertices/rosettes (>4 cells). For bicellular junctions, linear ROIs with a thickness of 5 pixels were drawn along the entire length of each junction. Background values were determined for each image by placing five circular ROIs with100-pixel diameter in a cell-free region (for Fig. 1A-C) or ROIs with 50-pixel diameter in a junction-free region (for Fig. 1D-F). The mean background intensity was subtracted from vertex and bicellular junction measurements. For each image, a length-weighted mean fluorescence intensity of bicellular junctions was calculated and used for normalization. Vertex intensities were then expressed relative to this bicellular reference (bicellular = 1). Data were visualized as bubble plots, in which the area of each bubble corresponded to the length of the traced bicellular junction, while tricellular and tetracellular/rosette vertices were plotted as normalized values with a uniform bubble size.

### Quantification of epithelial integrity of MDCK II cells

Fluorescence images of ZO-1 and DAPI from 4 coverslips for each cell clone or each treatment were used for quantification. For each coverslip, five non-overlapping fields of 1360 x 1024 pixels (corresponding to 216 x 163 µm) were collected, resulting in total of 20 images per cell clone or treatment. To assess epithelial integrity, ZO-1 staining was used to identify and manually mark all vertices in each image using ImageJ software. Vertices were classified as tricellular, tetracellular, or rosette-type (>5 cells), and the proportion of each category was calculated. Discontinuities in cell-cell junctions were also counted from ZO-1 images. Cell numbers were determined from the DAPI channel. Nuclei touching the image border were counted as 0.5 to correct for partial cells. The total number of vertices was divided by the number of nuclei to obtain the vertex-to-cell ratio. Similarly, the number of junctional discontinuities was normalized to the number of nuclei to yield the number of discontinuities per cell. Each image was treated as one sample (n = 20 per clone).

### Immunoblotting

Cells or bead-bound proteins were boiled in the SDS sample buffer for 5 min. Proteins were separated by SDS-PAGE using 5-20% gradient gels (Fujifilm Wako, Osaka, Japan), and transferred to a polyvinylidene fluoride (PVDF) membrane (Immobilon, Merck). The membranes were blocked with 5% non-fat dried milk in Tric-buffered saline (TBS) containing 0.1% Tween-20 (TBS-T) for 30 min at 20-28°C, followed by incubation with a primary antibody diluted in TBS-T overnight at 4°C. After washing with TBS-T, the membranes were incubated with an HRP-conjugated secondary antibody in TBS-T for 1 h at 20-28°C. After washing with TBS-T, the membranes were incubated with ECL Prime (GE Healthcare) and developed using LAS4000 (GE Healthcare). The images were processed using Photoshop.

### Co-immunoprecipitation

To examine the interaction between PAK4 and Afadin, HEK293T cells were co-transfected with the expression vectors using PEI-Max in a 6-well plate and cultured for 2 d. For FLAG-IP, cells were lysed with IP-lysis buffer (50 mM Tris-HCl, pH 7.4, 150 mM NaCl, 0.5% Nonidet P-40 [NP-40, #25223-04, Nacalai Tesque, Kyoto, Japan], 1 mM EDTA, 1 mM DTT, cOmplete protease inhibitor cocktail [#11836145001; Sigma-Aldrich]), precleared with Protein G-Sepharose (#17061801; Cytiva, Tokyo, Japan), and centrifuged at 15,000 x g for 20 min at 4°C. Supernatants were mixed with mouse-anti-DYKDDDDK mAb-bound Protein G-Sepharose and rotated for 2 hr at 4°C. Beads were washed five times using IP-wash buffer (50 mM Tris-HCl, pH 7.4, 150 mM NaCl, 0.05% NP-40, 1 mM EDTA, 1 mM DTT) and bead-bound proteins were eluted with SDS sample buffer. For GFP-trap, cells were lysed with GFP-trap lysis buffer (10 mM Tris-HCl, pH 7.5, 150 mM NaCl, 0.5% NP-40, 0.5 mM EDTA, cOmplete protease inhibitor cocktail) and left on-ice for 30 min with pipetting every 10 min. Lysates were centrifuged at 15,000 x g for 20 min at 4°C and diluted with 1.5x volume of dilution buffer (10 mM Tris-HCl, pH 7.5, 150 mM NaCl, 0.5 mM EDTA). Diluted lysates were mixed with GFP-trap magnetic agarose (#GTMA-20; Proteintech) and rotated for 1 hr at 4°C. Beads were washed five times with GFP-trap wash buffer (10 mM Tris-HCl, pH 7.5, 150 mM NaCl, 0.05% NP-40, 0.5 mM EDTA) using a magnet. Bead-bound proteins were eluted with SDS sample buffer.

### Transepithelial electrical resistance (TER) measurement

Cells were seeded onto 12-well polycarbonate Transwell filters with 0.4-µm pore size at 1.0 x 10^5^ cells/mL and cultured for 7 d. The transepithelial electrical resistance (TER) of the cell sheet was measured each day using a volt-ohm meter (Millicell ERS-2; EMD Millipore). All measurements were subtracted by a blank measurement of a Transwell filter with medium alone and then multiplied by the culture area of the Transwell filter to calculate the unit area resistance.

### AlphaFold3 prediction of the structure of PAK4-Afadin complex

The structures of the PAK4-Afadin complex, PAK4 alone, and Afadin alone were predicted using the AlphaFold server (https://alphafoldserver.com) (Abramson et al., 2024) and visualized using Open-Source PyMOL (The PyMOL Molecular Graphics System, Version 3.0 Schrödinger, LLC., Available at: http://www.pymol.org/pymol). PAE values were visualized with PAE viewer (https://pae-viewer.uni-goettingen.de) (Elfmann and Stülke, 2023). Possible interacting region was visualized with AlphaBridge (https://alpha-bridge.eu) (Álvarez-Salmoral et al., 2024).

### DNA constructs used in *Xenopus* experiments

*Xenopus* PAK4.L sequence (*X. laevis* v10.1) was synthesized by TWIST Bioscience (South San Francisco, CA, USA) and cloned into pCS2+/mNeon. Untagged *Xenopus* PAK4-NT was cloned by PCR amplifying the N-term fragment (amino acids: MFAKKKKRVEISAPSNFEHRVHTGFDQQEQKFTGLPRQWQSLIEESAKRPKPLVDPSYITTIKHVPQKTIVRGNKMSLDGSLAWLLDEFDDMSVCRSNS) from pCS2+/mNeon-PAK4 plasmid (described above) and inserting it into a pCS2+ vector digested with EcoR I and Xho I using Gibson cloning (Gibson et al., 2009) with the following primers: Forward 5′-CTTGTTCTTTTTGCAGGATCCCATCGATTCGAATTCACCATGTTTGCTAAGAAGAAGAAGC-3’; Reverse 5′-GTTTGCCGGTCGAACTCTTAACTCGAGCCTCTAGAACTATAGTGAGTCGTATT ACGT-3’. Constructs were verified by sequencing using Plasmidsaurus (Louisville, KY, USA). pCSf107mT/Lifeact-miRFP703 (Yamamoto et al., 2021) was a gift from Dr. Kazuhiro Aoki’s lab (Kyoto University, Japan). All other DNA constructs were previously described: pCS2+/TagBFP2-ZO-1 (Stephenson et al., 2019), pCS2+/PLEKHA7-mCherry (Higashi et al., 2019), and pCS2+/Angulin-1-3xGFP (Higashi et al., 2016).

### mRNA preparation

DNA was linearized using either Not I or Kpn I (in the case of ZO-1 construct). Linearized or non-linearized (in the case of Lifeact-miRFP703 construct) DNA was then transcribed using the mMessage mMachine SP6 Transcription Kit (#AM1340, Thermo Fisher Scientific). Transcribed mRNA was purified using the RNeasy Mini Kit (#74104, Qiagen, Germatown, MD, USA) and stored at -80°C until use.

### *Xenopus laevis* embryos and microinjections

All experiments conducted using *Xenopus laevis* embryos were carried out in compliance with the University of Michigan Institutional Animal Care and Use Committee and the U.S. Department of Health and Human Services Guide for the Care and Use of Laboratory Animals. The day prior to egg collection, female *Xenopus laevis* frogs (Xenopus 1 or National *Xenopus* Resource [NXR]) were induced to hyperovulate using human chorionic gonadotrophin (#198591, MP Biomedicals, Irvine CA, USA). The following day, eggs were collected from female frogs and fertilized *in vitro*. Fertilized embryos were dejellied approximately 30 min after fertilization and transferred to 0.1× MMR (10 mM NaCl, 0.2 mM KCl, 0.2 mM CaCl_2_, 0.1 mM MgSO_4_, and 0.5 mM HEPES, pH 7.4-7.6). Once embryos developed to the 2- or 4-cell stage, they were microinjected with the desired mRNAs. Cells were injected once per cell (for embryos at the 4-cell stage) or twice per cell (for embryos at the 2-cell stage). For PAK4, ZO-1, and PLEKHA7 colocalization data, each 5 nl of injection volume contained: 25 pg of PAK4-mNeon, 70 pg of TagBFP2-ZO-1, and 28 pg of PLEKHA7-mCherry. For PAK4 inhibitor experiments, each 5 nl of injection volume contained: 70 pg of TagBFP2-ZO-1, 25 pg of Angulin-1-3xGFP, and 150 pg of LifeAct-miRFP. For PAK4-NT experiments, each 5 nl of injection volume contained: 70 pg of TagBFP2-ZO-1, 150pg of LifeAct-miRFP, and 150-200pg of PAK4-NT. For ZnUMBA imaging, albino *Xenopus laevis* embryos were used. Embryos were injected with the following mRNA constructs: 70 pg of TagBFP2-ZO-1 and 150 pg of PAK4-NT, per 5 nl injection volume.

Injected embryos were then incubated in 0.1× MMR and allowed to develop for 16-24 h at 15°C or 13°C until they reached gastrula stage (Nieuwkoop and Faber stages 10-11).

### Zinc-based Ultrasensitive Microscopic Barrier Assay (ZnUMBA)

For ZnUMBA experiments, gastrula-stage embryos were microinjected one time in the blastocoel with 10 nl of a mixture of 100 μM CaCl_2_, 100 μM EDTA, and 1 mM FluoZin-3 (#F24193, Thermo Fisher Scientific). FluoZin-3–injected embryos were allowed to recover from microinjection for 5-10 min in 0.1× MMR. After recovery, embryos were mounted in 1 mM ZnCl_2_ in 0.1× MMR before imaging (Higashi et al., 2023b).

### Immunofluorescence staining of *Xenopus* embryos

PFA fixation was performed as follows: gastrula-stage albino embryos were placed in a solution of 1.5% PFA, 0.25% glutaraldehyde, 0.2% Triton X-100, and Alexa Fluor 568 phalloidin (#A12380, Thermo Fisher Scientific) (1:100) in 0.88× MT buffer (80 mM K-PIPES, 5 mM EGTA, 1mM MgCl_2_, pH 6.8) and allowed to fix on a shaker overnight at room temperature. Fixed embryos were quenched in 100 mM sodium borohydride in PBS on a shaker for 1 hr at room temperature. Embryos were then bisected to keep the animal cap and blocked with blocking solution (10% FBS, 5% DMSO, 0.1% NP-40 in 1× TBS) overnight on a nutator at 4°C. The animal caps were then incubated overnight on a nutator at 4°C in the blocking solution with mouse anti-PAK4 mAb (#sc-390507, Santa Cruz Biotechnology) (1:50). Next, they were washed three times in blocking solution and incubated overnight on a nutator at 4°C in the blocking solution with goat Alexa Fluor 488 anti-mouse pAb (A-11001, Invitrogen) (1:200), and Alexa Fluor 568 phalloidin (A12380, Invitrogen) (1:100). The next day, embryos were washed in 0.1% NP-40 in TBS (TBSN), then incubated in 1:1000 DAPI (10 mg/ml stock) diluted in TBSN for 30 min at room temperature on the nutator. Next, they were washed for 10 min with TBSN five times. Finally, embryos were washed and mounted in PBS before imaging.

### Live imaging of *Xenopus* embryos

Live imaging videos of gastrula-stage *Xenopus* embryos were acquired using an Olympus FluoView 3000, and 60X supercorrected Plan Apo N 60X OSC objective (NA = 1.4, working distance = 0.12 mm), with mFV10-ASW software. Embryos were mounted as described previously (Reyes et al., 2014). Embryos were imaged with a scanning speed of 2 or 4 µs/pixel, a 512 × 512 field of view, 1× zoom, capturing 6-10 of the most apical slices of the animal cap of the embryo, with a step size of 0.5 µm. For live imaging of PAK4 with ZO-1 and PLEKHA7 (Fig. 2E), embryos were imaged with a scanning speed of 4 µs/pixel, a 512 × 512 field of view, 1× zoom, capturing 50 of the most apical optical slices of the animal cap of the embryo, with a step size of 0.5 µm (Higashi et al., 2019). Fixed embryos were imaged with a scanning speed of 4 µs/pixel, a 512 × 512 field of view, 1× zoom, capturing 40 optical slices of the embryo, with a step size of 0.5 µm.

### Drug treatment of *Xenopus* embryos

The PAK4 inhibitor, PF-3758309 (S7094, Selleck, Houston, TX, USA), was resuspended in DMSO as a 10 mM stock solution, aliquoted and stored in -80°C.

Following mRNA injection at the 2- or 4-cell stages, embryos were incubated overnight at 15°C in either 0.1% DMSO (Control) or 10 µM PAK4 inhibitor in 0.1× MMR.

### Image analysis of *Xenopus* embryos

#### Quantification of the localization of PAK4 at cell-cell junctions

Gastrula-stage embryos expressing PAK4-mNeon together with PLEKHA7-mCherry and TagBFP2-ZO-1 were live imaged as described above. Square images (50 pixels × 50 pixels) of straight junctions were excised from the original images, and 50 z-slices across the junctions were averaged using ImageJ. Fluorescence intensity profiles along x- and z-axes were determined by multiple line scans using ImageJ (four line scans from each embryo) (Higashi et al., 2019). This process was repeated for 16 junctions from 5 embryos and aligned using the highest ZO-1 intensity; mean and SD was calculated. For measuring the intensity of PAK4 and ZO-1 at the vertices, a circular ROI of size 10 x 10 pixels was used to measure the intensity at 16 bicellular junction (BCJ), tricellular vertices (TCJ) or higher-order vertices (4-way and rosettes) from maximum z-projected images of 4 different embryos using Fiji. The intensity at TCJs and higher-order vertices were normalized by their counterpart BCJ intensity.

#### Quantification of PAK4 intensity at remodeling vertices

For measuring the change in PAK4 intensity, a circular ROI of size 3 x 3 pixels (1.24 µm x 1.24 µm) was used to measure the intensity at remodeling vertices, non-remodeling vertices, and the cytoplasm over 60-90 mins using Fiji. The intensity was normalized as follows: ((Intensity at vertex - Cytoplasmic Intensity) / Cytoplasmic Intensity). To measure the change in angle, the angle tool in Fiji was used with the lines spanning 5 µm along each junction. The early stage of remodeling was defined by time point when the vertex starts actively moving. The late stage is towards the end as the vertex stabilizes. The middle stage is the time point exactly halfway. A non-remodeling vertex was chosen in the same field of view as a remodeling vertex and the measurements were taken for all the timepoints as its remodeling vertex counterpart. PAK4 intensity at the remodeling vertices at early, middle, and late stages was normalized as follows:(Intensity at remodeling vertex/Average intensity of non-remodeling vertices at that stage).

#### Quantification of cell packing upon PAK4 inhibitor treatment

Distribution of cell-cell interfaces after cytokinesis: Using cells expressing TagBFP2-ZO-1, the distribution of Type I (daughter-daughter interface), Type II (4-way junction), and Type III (neighbor-neighbor interface) interfaces was determined for each embryo. Movies of both control and PAK4-inhibited embryos were analyzed from 13 to 60 minutes after ingression began. Rosette resolution: Using cells expressing TagBFP2-ZO-1, the number of rosettes (where 5 or more cells meet) at the start of imaging was determined for each embryo. The marked rosettes were then analyzed for both control and PAK4 inhibited embryos for 60-80 minutes. The time when the 5-way vertex starts to resolve from the beginning of the movie was noted as the time of rosette resolution.

#### Quantification of cytokinesis failure

Using cells expressing TagBFP2-ZO-1, cell division events were followed for 13-96 minutes after ingression began in 5 controls (46 cell divisions) and 8 PAK4-NT-expressing embryos (32 cell divisions). Formation of a 4-way vertex with clear ZO-1 intensity was marked as cytokinesis completion, whereas dissolution of the cytokinetic ring, inability to form a 4-way vertex with clear ZO-1 intensity, or breaking off of an already formed daughter-daughter interface were considered cytokinetic failure.

#### ZnUMBA

Whole field intensity analysis: All movies were imaged using the same imaging parameters. The images were maximum z-projected and contrast adjusted to 0-4095 in Fiji. Whole field intensity was measured by Fiji at 0 min, ∼7 min, and ∼15 min since the start of imaging. If the junctions were not visible at 15 mins in the FluoZin3 channel, it was considered a global barrier leak.

Junctional intensity analysis: Cell-cell junctions were traced using the TagBFP2-ZO-1 channel and threshold option in Fiji. Post thresholding, the image was made binary and an ROI was created tracing the cell-cell junction. The ROI was superimposed on the FluoZin3 channel, and the intensity was measured in Control, PAK4 inhibitor-treated embryos, and PAK4-NT-expressing embryos.

Quantification of leaky vertices: Barrier leaks were counted manually by identifying vertices where FluoZin3 intensity increased over baseline intensity, as revealed by a change in color using the FIRE LUT applied in ImageJ. This count was then normalized by the number of vertices present in the field of view, and the percentage of leaky vertices was calculated. In the case of PAK4-NT quantification, embryos with global barrier leaks were excluded from this analysis.

Quantification of FluoZin3 intensity at higher-order vertices: A circular ROI of size 7 x 7 pixels (2.90 µm x 2.90 µm) was used to measure intensity at higher-order vertices (4-way or higher) and in the cytoplasm at 0 min, ∼7min, and ∼15 min using Fiji, and the intensity was normalized as follows: (Intensity at higher-order vertex - Cytoplasmic Intensity) / (Cytoplasmic Intensity).

## Video editing

Videos were initially generated and concatenated using FiJi 2.16.0/1.5p. Overlay animations were added using Adobe Premiere pro 25.5.

## Statistical analysis and plotting

Data shown in Figs. 1B-C, 1E-F, 3B-D, 5B-D, 6B-D, 6G-I, S2B-D, and S2G-I are presented as the mean ± standard error (S.E.M.) using Microsoft Excel 365. Data in Figs 8A-F and 8J-K are presented as the mean ± standard deviation using Microsoft Excel 365. Statistical significance of the differences was evaluated using a two-tailed unpaired Welch’s *t*-test (Figs. 1B-C, 1E-F, 3B-D, 5B-D, 6B-D, 6G-I, S2B-D, S2G-I). Bonferroni correction was used to address the multiple comparison problem in Figs. 3B-D, 5B-D, 6B-D, 6-I, S2B-D, S2-I. All tests were performed using Microsoft Excel 365. Weighted average intensity at bicellular junctions in Figs. 1B and 1E was calculated using R software (https://www.r-project.org) using weighted.mean() function.

For data presented in Figs. 2, 4, 7, 9, 10, and S5, all graphs were plotted using python 3.8.10 using the following packages: matplotlib (3.7.5), numpy (1.24.4), openpyxl (3.1.5), pandas (2.0.3), pip (25.0.1), scipy (1.10.1), seaborn (0.13.2), statsmodels (0.14.1), and statannotations (0.7.2). Some of the basic data handling was performed on Microsoft Excel 365. A two-tailed unpaired Student’s *t*-test was performed for Figs. 2C, 2D, 2G, 4G, 7C, 9D, 10C-D, 10F, and S5B-C. A two-tailed unpaired Welch’s *t*-test was used for Figs. 9C and 10E.

## Acknowledgements

We would like to express our gratitude to Misa Satoh, Seiko Watanabe, Keiko Watari, Chiaki Ozaki, and Joji Kai for their technical assistance, Ryutaro Shirakawa (Japan Institute for Health Security) for the critical review of our manuscript, and Janis F. Miroll for assistance with statistical analysis and plotting with Python. This work was funded by the JSPS (Japan Society for the Promotion of Science) KAKENHI (Grant Number 24K09456), the 2024 Naito Grant for the advancement of natural science from Naito Foundation, the 2024 Narishige Zoological Science Award from Narishige Fund to T.H., and NIH R35GM153204 to A.L.M. The authors declare no competing financial interests.

## Author contributions

This project was designed by T.H. All the experiments and data analyses using MDCK II cells were performed by T.H. with input from H.C. All the experiments and data analyses using *Xenopus* embryos were performed by B.A. A.C. and A.Y.H made DNA constructs and performed immunostaining of mouse tissues, respectively. A.L.M. and T.H. supervised the project. A.L.M. and T.H. were responsible for project administration. B.A. and T.H. wrote the manuscript with input from other authors, and all authors reviewed and revised the manuscript.

## Abbreviations

AJ: adherens junction
PAK4: p21-activated kinase 4
PBD: p21-binding domain
TJ: tight junction
tAJ: tricellular adherens junction
tTJ: tricellular tight junction
TER: transepithelial electrical resistance
ZnUMBA: Zinc-Based Ultrasensitive Microscopic Barrier Assay

## Video legends

**Video 1. Time-lapse confocal imaging of gastrula-stage *Xenopus* embryos showing increased PAK4 localization at remodeling vertex.** Video shows PAK4 (PAK4-mNeon, green Fire Blue LUT) intensity at a non-remodeling vertex (left) and remodeling vertex (right). Yellow arrows show PAK4 accumulation during remodeling, while red arrows point out the non-remodeling vertex as well as the end of remodeling. Time interval, 24 sec; video frame rate, 10 fps. Related to Fig. 2F.

**Video 2. Time-lapse confocal imaging of gastrula-stage *Xenopus* embryo showing impaired higher-order vertex resolution remodeling upon PAK4 inhibition.** Video illustrates that the PAK4 inhibitor treated embryo (right) has higher number of rosettes (yellow circules) and these take longer to resolve compared to control (DMSO treated) embryo (left). Time interval, 34 sec (left video), 32 sec (right video); video frame rate, 15 fps. Related to Fig. 4.

**Video 3. Time-lapse confocal imaging showing cytokinetic failure and junctional discontinuities in PAK4-NT-expressing gastrula-stage *Xenopus* embryos.** First video illustrates that inhibiting the recruitment of endogenous PAK4 via expressing PAK4-NT causes cytokinetic failure. The cell starts dividing marked by cytokinetic furrow ingression (white arrow). Eventually the daughter-daughter cell interface breaks (cyan arrow) leading to cytokinesis failure. Time interval, 37 sec; video frame rate, 7 fps. Related to Figure 7D, E. Second video illustrates junction discontinues in PAK4-NT embryos. The yellow arrowheads show some of the breaks and discontinues in the junctions. Time interval, 37 sec; video frame rate, 7 fps. Related to Figure 7F.

**Video 4. Time-lapse confocal imaging of gastrula-stage *Xenopus* embryos showing barrier leaks upon PAK4 inhibition.** PAK4 inhibitor-treated embryos (right) have increased baseline FluoZin3 intensity (FIRE LUT) and an increased frequency of tight junction leaks compared with controls (left). Time interval, 2:44 (min:sec); video frame rate, 3 fps. Brightness and contrast were adjusted to the same levels for both videos. Related to Figure 9B.

**Video 5. Time-lapse confocal imaging of gastrula-stage *Xenopus* embryos showing barrier leaks upon PAK4-NT expression.** PAK4 NT-expressing embryos (right) have increased baseline FluoZin3 intensity (FIRE LUT), global barrier disruption (#2), and an increased frequency of tight junction leaks at tricellular vertices (#1) compared with controls (left). Time interval, 2:44 (min:sec); video frame rate, 3 fps. Brightness and contrast were adjusted to the same levels for all the videos. Related to Figure 10A.

