## Supplemental Figures for "PAK4 promotes vertex remodeling to maintain epithelial integrity and barrier function"

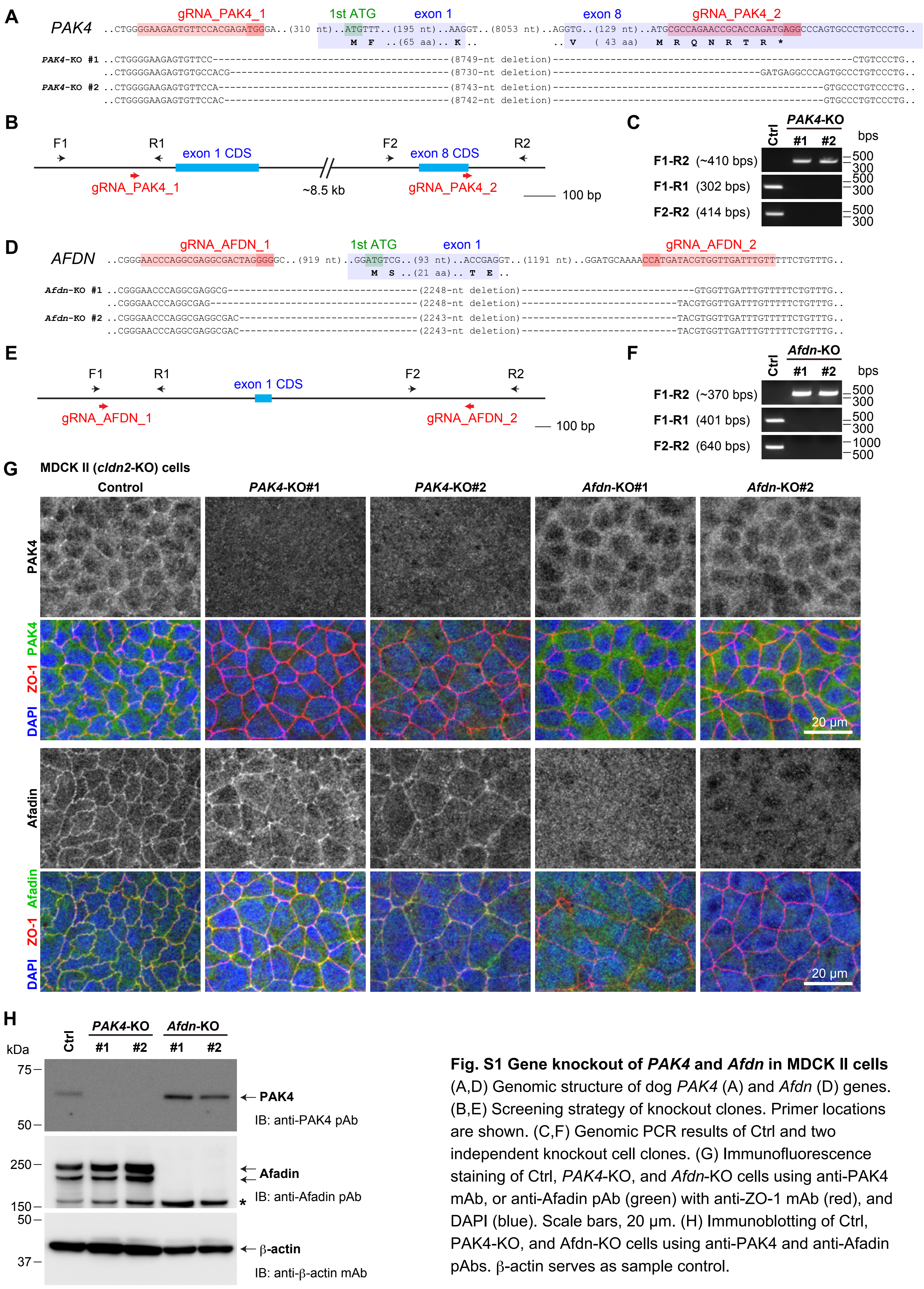


**Fig. S1 Gene knockout of *PAK4* and *Afdn* in MDCK II cells**
(A,D) Genomic structure of dog *PAK4* (A) and *Afdn* (D) genes. (B,E) Screening strategy of knockout clones. Primer locations are shown. (C,F) Genomic PCR results of Ctrl and two independent knockout cell clones. (G) Immunofluorescence staining of Ctrl, *PAK4*-KO, and *Afdn*-KO cells using anti-PAK4 mAb, or anti-Afadin pAb (green) with anti-ZO-1 mAb (red), and DAPI (blue). Scale bars, 20 μm. (H) Immunoblotting of Ctrl, PAK4-KO, and Afdn-KO cells using anti-PAK4 and anti-Afadin pAbs. β-actin serves as sample control.

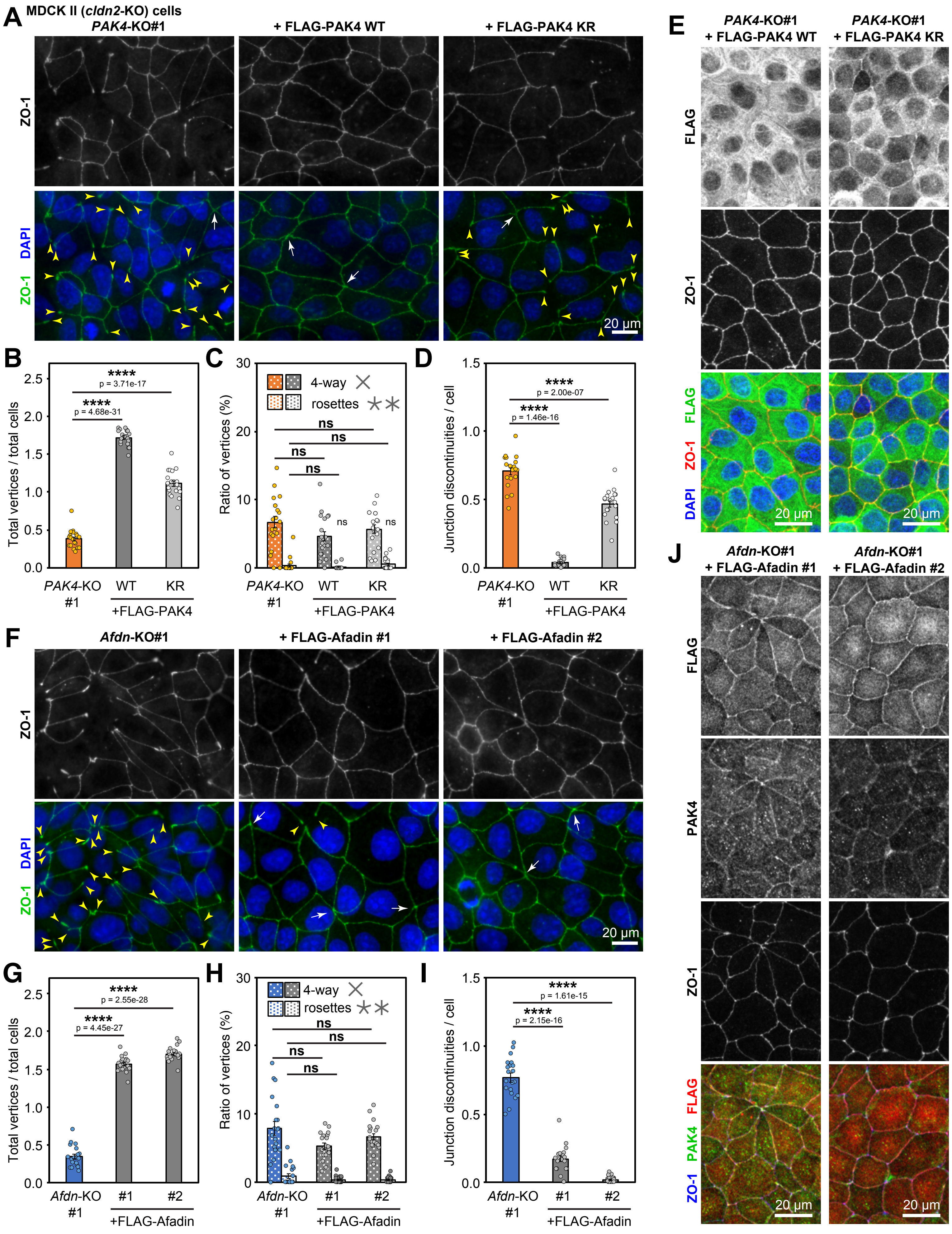


**Fig. S2 Rescue of epithelial integrity phenotypes of *PAK4*-KO and *Afdn*-KO cells**

(A,E,F,J) Immunostaining of *PAK4*-KO cells, *PAK4*-KO cells expressing FLAG-PAK4 WT or KR, *Afdn*-KO cells, *Afdn*-KO cells expressing FLAG-Afadin cultured on glass coverslips for 40 hr (A,F) or 48 hr (E,J) using anti-ZO-1 mAb and DAPI (A,F) with anti-FLAG mAb (E) or anti-FLAG mAb and anti-PAK4 mAb (J). (B-D,G-I) Quantification of the vertex-to-cell ratio (B,G), proportions of 4-way vertices and rosettes (C,H), and number of junction discontinuities (D,I). Data are compared with a two-tailed Welch's *t*-test with Bonferroni's correction (ns, $p > 0.05$; ****, $p < 0.0001$). S.E.M (bars) and individual values (circles) are shown.

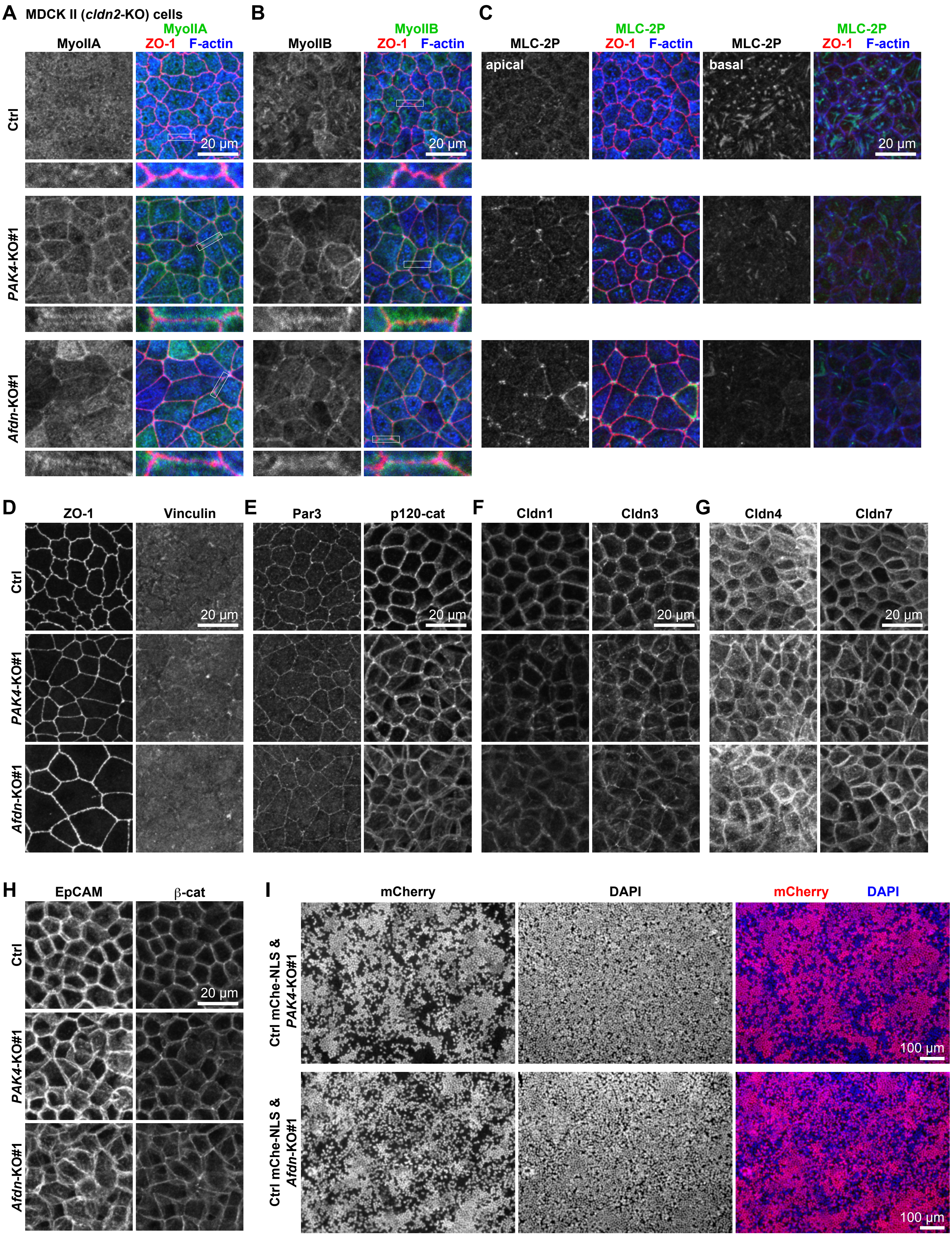


**Fig. S3 Characterization of *PAK4*-KO and *Afdn*-KO cells**

(A-H) Immunostaining of *PAK4*-KO and *Afdn*-KO cells cultured on Transwell filters for 7 days using antibodies against indicated proteins. Scale bars, 20 µm. (I) Mixed culture of *PAK4*-KO or *Afdn*-KO cells with Ctrl mChe-NLS cells were stained with DAPI (blue). Note that KO cells (cells without mCherry signal [red]) constitute almost half of the cell population, suggesting that there is no or little, if any, competitive removal of cell clones. Scale bars, 100 µm.

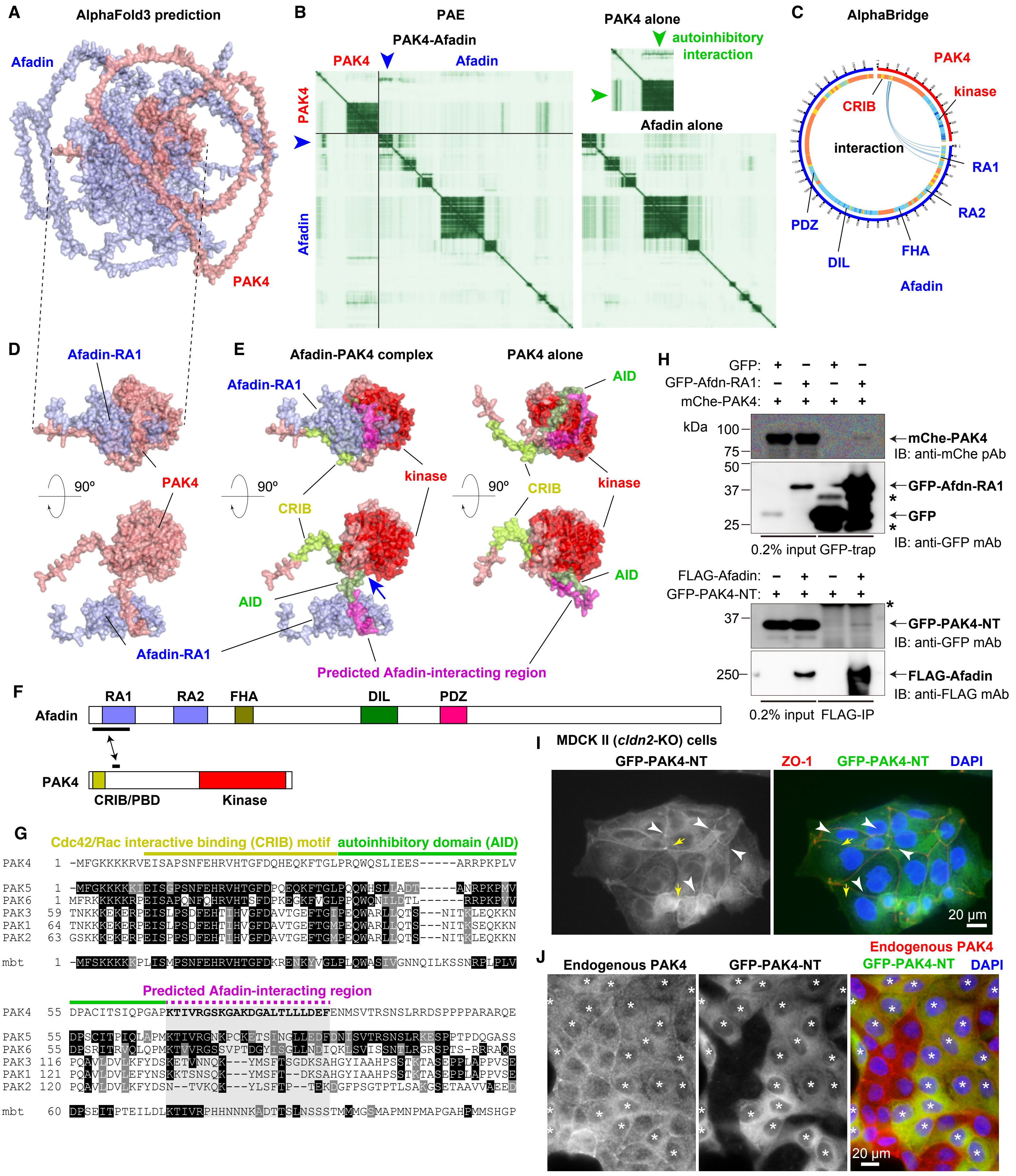


**Fig. S4 Structural prediction of PAK4-Afadin complex and PAK4-NT construction**

(A-E) 3D structure (A,D,E), PAE map (B), and AlphaBridge map (C) of PAK4-Afadin complex predicted by AlphaFold3. (F) Domain organization of Afadin and PAK4. RA, ras association; FHA, folkhead-associated; DIL, dilute; PDZ, PSD95-Dlg-ZO1; CRIB, Cdc42/Rac interactive binding; PBD, p21-binding domain. (G) Alignment of PAK4 sequence. (H) Coimmunoprecipitation of mChe-PAK4 and GFP-Afdn-RA1 or GFP-PAK4-NT and FLAG-Afadin from HEK293T cells. (I) Immunostaining of Ctrl MDCK II cells expressing GFP-PAK4-NT (green) using anti-ZO-1 mAb (red) and DAPI (blue). Scale bar, 20 µm. (J) Immunostaining of mixed culture of Ctrl MDCK II cells and GFP-PAK4-NT-expressing cells using anti-PAK4 mAb (red) and DAPI (blue). Note that cells expressing GFP-PAK4-NT (asterisks) lose endogenous PAK4 localization at cell-cell junctions. Scale bar, 20 µm.

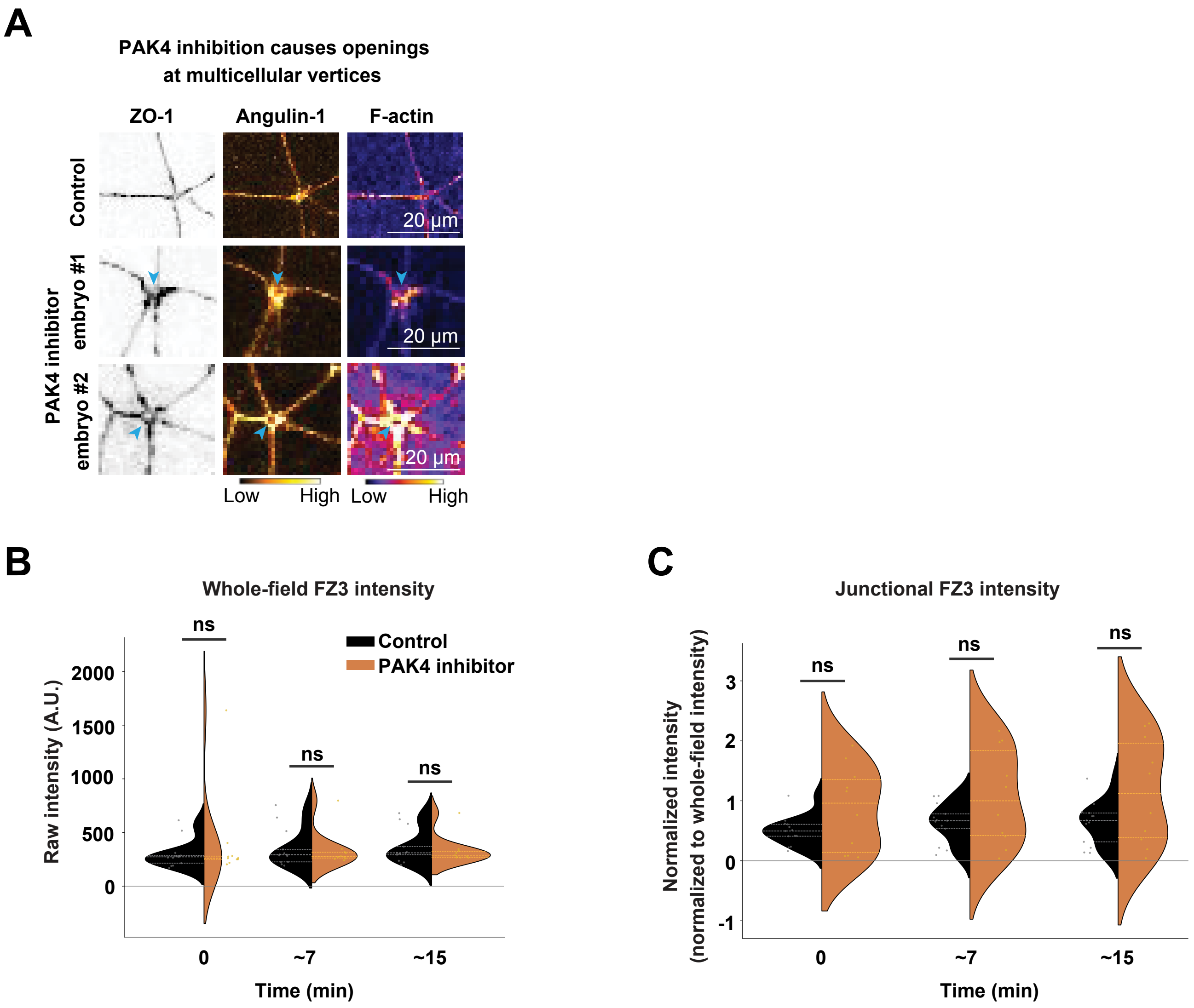


**Fig. S5 PAK4 inihibition causes openings at cell vertices**
(A) Live confocal images of epithelial cells in the animal hemisphere of gastrula-stage *Xenopus* embryos expressing TagBFP-ZO-1, Angulin-1-3xGFP, and LifeAct-miRFP703 treated with DMSO or 10 µM PAK4 inhibitor. Blue arrowheads indicate the opening at multicellular vertex in PAK4 inhibitor-treated embryos. Scale bars, 20 µm. (B-C) ZnUMBA assay of DMSO- or PAK4 inhibitor-treated embryos. Quantification of whole-field (B) and junctional (C) FluoZin-3 intensities at 0, ~7, and ~15 min after the onset of imaging. ns, $p > 0.05$. (control: 13 embryos; PAK4 inhibitor: 10 embryos; 3 independent experiments [B]; control: 13 embryos; PAK4 inhibitor: 9 embryos; 3 independent experiments [C])
